# Layered social competition coordinates reproductive hierarchy formation in ants

**DOI:** 10.64898/2026.05.11.724417

**Authors:** Shao Zhi George Liu, Manon Ben Kiran, Yu-Chin Lin, Rosie Wang, Yuwei Zhong, Yen-Chung Chen, Muhammad Zain Naveed, Claude Desplan, Ching-Han Lee

## Abstract

Social interactions establish reproductive hierarchies and regulate division of labor in eusocial insect colonies. In the ant *Harpegnathos saltator*, queen loss induces a subset of workers to transition into reproductive pseudo-queens, known as gamergates, through a ritualized antennal-dueling tournament that can last for more than a month. Gamergate transition is associated with sustained dueling engagement, yet the interaction networks and social dynamics that shape reproductive hierarchy formation remain unclear. Here, we combine long-term automated tracking with behavioral classifiers to systematically quantify social interactions, including antennal dueling and two distinct biting behaviors: aggressive grooming and mandible locking. We find that antennal dueling is organized into two activity states, an engaged state and a disengaged state, and that persistent maintenance of the engaged state, rather than transient entry into it, predicts successful transition to gamergate fate. Early access to abundant food shapes dueling activity and biases ants toward entry into the engaged state. Moreover, engaged-state individuals exhibit a strong assortative preference for dueling with one another and cluster spatially within the colony. We further show that mandible locking precedes transitions out of the engaged state, whereas aggressive grooming reinforces the disengaged state, together consistent with a role in stabilizing worker fate. Our findings reveal how structured social interactions organize reproductive hierarchy formation by regulating stable behavioral states associated with caste fate.

## Introduction

Across group-living animals, dominance hierarchies arise through diverse forms of social interaction and can regulate access to reproduction ^1–3^. Eusocial insects exhibit an extreme form of reproductive hierarchy, in which the queen caste is specialized for reproduction, whereas the worker caste performs most colony tasks while maintaining reduced or largely inactive ovaries. This caste differentiation is typically established during developmental transitions, in which nutritional, endocrine, and gene regulatory mechanisms canalize individuals into reproductive queens or non-reproductive workers ^4,5^. After caste identity is established, reproductive individuals can further regulate worker physiology and behavior through social interactions such as pheromonal communication and aggression ^6,7^, while social and environmental cues, such as brood care, also reinforce caste-specific behavioral states ^8,9^.

Caste identity is not always fixed in adulthood. In some ant species, adult workers retain reproductive capacity and can transition into reproductive individuals, known as gamergates, in response to queen loss and reproductive vacancy ^10^. This plasticity can enhance colony fitness by rapidly restoring reproduction after the accidental death or removal of the queen. Gamergates are found in multiple ant lineages, predominantly within Ponerinae but also in Myrmicinae and Ectatomminae, and reproductive hierarchy is re-established through species-specific social interactions ^11,12^. In *Diacamma ceylonense*, all workers are eclosed with similar reproductive potential, but the resident gamergate mutilates the workers’ gemmae, small thoracic appendages required for mating, thereby enforcing reproductive hierarchy ^13^. In *Dinoponera quadriceps*, reproductive status is regulated through dominance interactions among workers and reinforced by chemical signaling that can trigger punishment of challengers ^14,15^. Although gamergate emergence has been characterized in several ant species, how dynamic social interactions regulate hierarchy formation among reproductively plastic workers remains unclear.

The Indian jumping ant, *Harpegnathos saltator*, provides an exceptional system for dissecting adult caste plasticity ^16^. In colonies with reproductive individuals (queens or gamergates), worker reproduction is suppressed by queen-derived pheromones and reinforced by worker policing ^17^. Upon the loss of the queen, workers enter a ritualized dueling tournament characterized by rapid antennal strikes through which reproductive hierarchy is re-established ^16,18,19^. Previous studies have shown that the worker-to-gamergate transition is accompanied by extensive neurohormonal and molecular remodeling ^16,20,21^. In workers, juvenile hormone and corazonin promote worker-like physiology and behavior ^22^, whereas in gamergates, ecdysone-related signaling and nutrient-sensing pathways (notably insulin/IGF signaling) support reproduction and longevity ^23^. Neurotransmitter profiles also shift during caste transition, with elevated dopamine emerging as an important regulator of reproductive activation and caste-associated behavioral change ^21^. In addition, socially regulated hormones act through Krüppel homolog-1 (Kr-h1) to maintain caste-specific brain states by suppressing inappropriate transcriptional and behavioral programs ^24^. Despite advances in characterizing the molecular mechanisms that distinguish workers and gamergates, the full behavioral dynamics underlying social hierarchy formation remain poorly understood. Not all individuals that engage in antennal dueling ultimately become gamergates ^16^, and the factors that determine why some ants persist whereas others withdraw from the tournament remain elusive. Previous studies have largely relied on manual behavioral annotation within restricted time windows, which may overlook the fine temporal structure of social interactions during hierarchy formation. Manual annotation is also poorly suited to resolving the full interaction dynamic underlying hierarchy formation, because antennal dueling is highly frequent and long-lasting and involves complex, repeated exchanges among many individuals. Moreover, secondary behaviors such as biting and grooming have likewise rarely been quantified systematically. Thus, a more comprehensive behavioral measurement framework is needed to reconstruct the social interaction history that shapes individual caste fate.

SLEAP (Social LEAP Estimates Animal Poses) is a deep learning-based framework for multi-animal pose estimation that tracks user-defined body keypoints over time, such as the head, thorax, abdomen, and appendages ^25^. To enable long-term, high-resolution behavioral analysis with persistent individual identity, we used NAPS (ArUco Plus SLEAP), a hybrid framework that integrates SLEAP-based pose estimation with ArUco fiducial markers, which encode unique IDs for long-term individual identification ^26^. We further developed neural-network classifiers to quantify three social behaviors during caste transition: antennal dueling, aggressive grooming, and mandible locking. Within the antennal dueling tournament, we identified three caste-fate trajectories: High-dueling ants that ultimately become gamergates, Low-dueling ants that remain workers, and a third group that engages intensely in dueling but later withdraw and remain workers. Based on these trajectories, we divided the tournament into three stereotyped phases: adaptation, engagement, and stabilization. The other two social behaviors contributed to hierarchy formation in distinct ways: mandible locking preceded transitions out of the engaged state, whereas aggressive grooming is enriched after withdrawal and may stabilized the disengaged state. Together, we established automated behavioral quantification as a powerful approach for resolving the fine-scale social dynamics that govern hierarchy formation and caste fate during adult plasticity.

## Results

### Automated quantification of antennal dueling reveals a three-phase tournament

To quantify ant behavior in an unbiased and high-throughput manner, we adopted NAPS, a machine-learning tracking system that combines ArUco identity tags with the SLEAP posture framework to track individual identities alongside defined anatomical keypoints (e.g., antennae, head, thorax and abdomen) across all frames ^25–27^ (see Materials and Methods). This automated pipeline enabled identity conserved pose-tracking of individual ants over long recording periods **(Figure 1A, <u>Video S1</u>)**. To investigate social behaviors during caste transition in *H. saltator*, we established four queenless “transition colonies”, each with 15 age-matched young workers (∼14 days post-eclosion). Recording began the day after setup and continued for 28 days, 12 h per day during the light phase. We then applied NAPS to these recordings and assigned caste fate (gamergate vs. worker) at the end of recording by assessing ovarian development and caste-specific brain markers (See Materials and Methods; **Table 1**). In addition, we measured head width for each ant as a proxy for body size and found that gamergates had slightly larger head widths than workers (**Figure S1C**; 2.3% larger, p = 0.012).

**Figure 1.**
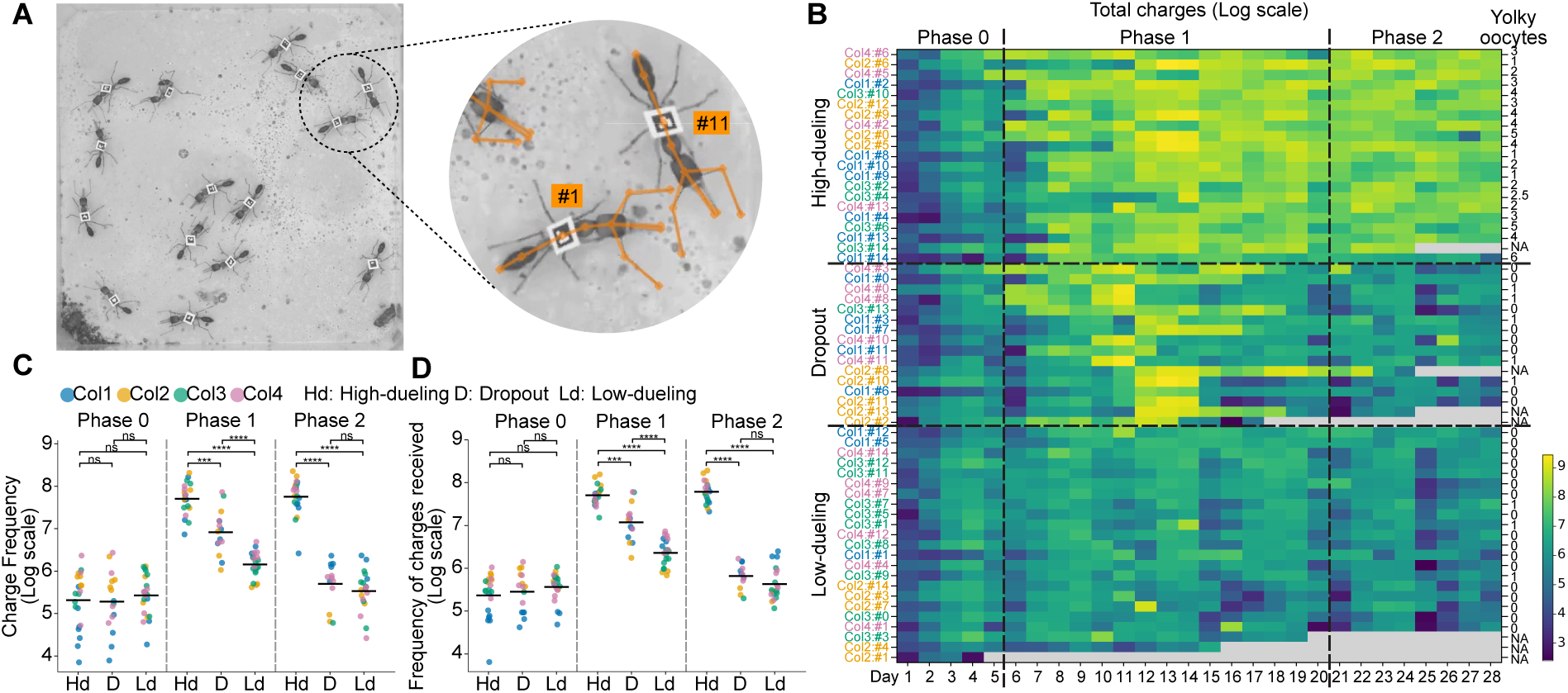
Automated high-throughput tracking reveals a three-phase antennal dueling tournament with three caste-fate trajectories. (**A**) A transition colony of 15 age-matched workers tagged with ArUco tags. Inset shows the SLEAP skeletal key point overlay. (**B**) Pooled heatmap of total daily charging events (natural log scale) across 28 days for four colonies. Rows represent individual ants, grouped by caste-fate trajectory and sorted by mean Phase 1 intensity. The rightmost column shows yolky oocyte counts. Gray cells indicate missing recording days because of ant death (NA). (**C-D**) Phase-stratified dot plots of natural-log-scaled total charge frequency (**C**) and total charges received frequency (**D**). Each dot represents the phase-averaged charge frequency for one ant, colored by colony (1-4). Horizontal bars indicate colony-pooled means. Significance brackets indicate p values from colony-blocked permutation tests: ns, not significant; **, p < 0.01; ***, p < 0.001; ****, p < 0.0001.

We first used this keypoint dataset to detect and quantify the antennal dueling tournament, which was considered strongly associated with a worker’s transition to a reproductive gamergate fate ^16^. Antennal dueling is a stereotyped sequence in which two workers charge toward one another and exchange rapid antennal strikes. During each charge, ants lean their bodies forward and strike the opponent with their antennae, most often toward the target’s head **(Figure S1B)**. To automatically detect individual charging events, we trained a spatiotemporal graph convolutional network (ST-GCN) ^28^, hereafter referred to as Duel**-**Tracker, on thousands of manually annotated clips to learn the spatiotemporal pose dynamics of the charger and target (see Materials and Methods). We used the model to quantify total charging events for each ant over the 28-day recording **(Figure 1B)**. The model captured up to 11,448 such charging events per day, per ant, demonstrating a throughput and resolution not achievable by manual quantification. We recapitulated previous findings that gamergate-fated ants exhibit higher charging frequencies than worker-fated ants and sustained intensive dueling for more than a month ^21^ **(Figure 1B)**. Additionally, we detected an average of ∼500 charging events per day among worker-fated ants, and ∼300 even late in the transition, challenging the view that workers cease dueling soon after tournament onset **(Figure 1B-C)** ^21^. We found that charging followed a highly structured temporal pattern: activity remained low during the first 5-6 days, increased over the following 15 days, and ultimately stabilized, with sustained intense charging largely confined to ants that later became gamergates. We therefore divided the tournament dynamics into three phases: Phase 0 (adaptation), Phase 1 (engagement), and Phase 2 (stabilization) **(Figure 1B-C)**. Based on these phases, we defined three caste-fate trajectories: High-dueling ants exhibited intense charging in Phase 1 and sustained it in Phase 2, ultimately becoming gamergates, whereas Low-dueling ants maintained low charging throughout all three phases and remained workers. Intriguingly, about one-fourth of the ants (16 out of 60) showed intense charging in Phase 1 but dropped out in Phase 2 and ultimately remained workers; hereafter, we refer these ants as Dropouts **(Figure 1B-C)**.

A key component of our dueling model was target assignment. For each charging event, we defined the target as the individual whose body keypoints were most frequently closest to the charging ant’s mandibles during the 2-second interaction window. We then quantified the number of charges each ant received during the caste transition **(Figure 1D)**. The temporal pattern of charges received closely mirrored that of charges given: High-dueling ants and Dropouts received intense charges in Phase 1, indicating active engagement in the tournament, whereas only High-dueling ants continued to receive intense charges in Phase 2, indicating that future gamergates primarily duel with one another **(Figure 1D)**.

### Future gamergates exhibit high responsiveness and prolonged reciprocal dueling

We next asked whether individual charging events are organized into larger interaction structures during antennal dueling. Antennal dueling comprises back-and-forth charges between opponents. Early in the tournament, charges were frequently non-reciprocal, with a focal ant charging a target that did not respond **(<u>Video S2</u>)**. Later, as dueling intensified, exchanges became reciprocal, with opponents charging back and forth from 2 up to 36 charges **(<u>Video S3</u>)**. To dissect this, we classified charges as initiating or response events and calculated the response rate **(Figure 2A-B and S2A-C)**. We defined a response charge as any charge by a focal ant that had received a charge from its target within the preceding 2-second window, reflecting a reactive response rather than an initial decision to engage. All other charges were classified as initiating charges. We next defined the response rate as the probability that an ant charged back within 2 seconds of receiving a charge. We found that High-dueling ants and Dropouts directed and received more initiating charges than Low-dueling ants in Phase 1 **(Figure 2A and S2A-B)**. In addition, High-dueling ants and Dropouts showed higher response rates in Phase 1 **(Figure 2B and S2C)**, indicating active engagement in the tournament. In Phase 2, only High-dueling ants maintained elevated initiating charges and response rates **(Figure 2A-B and S2A-C)**, consistent with the conclusion that ants sustaining dueling activity ultimately became gamergates.

**Figure 2.**
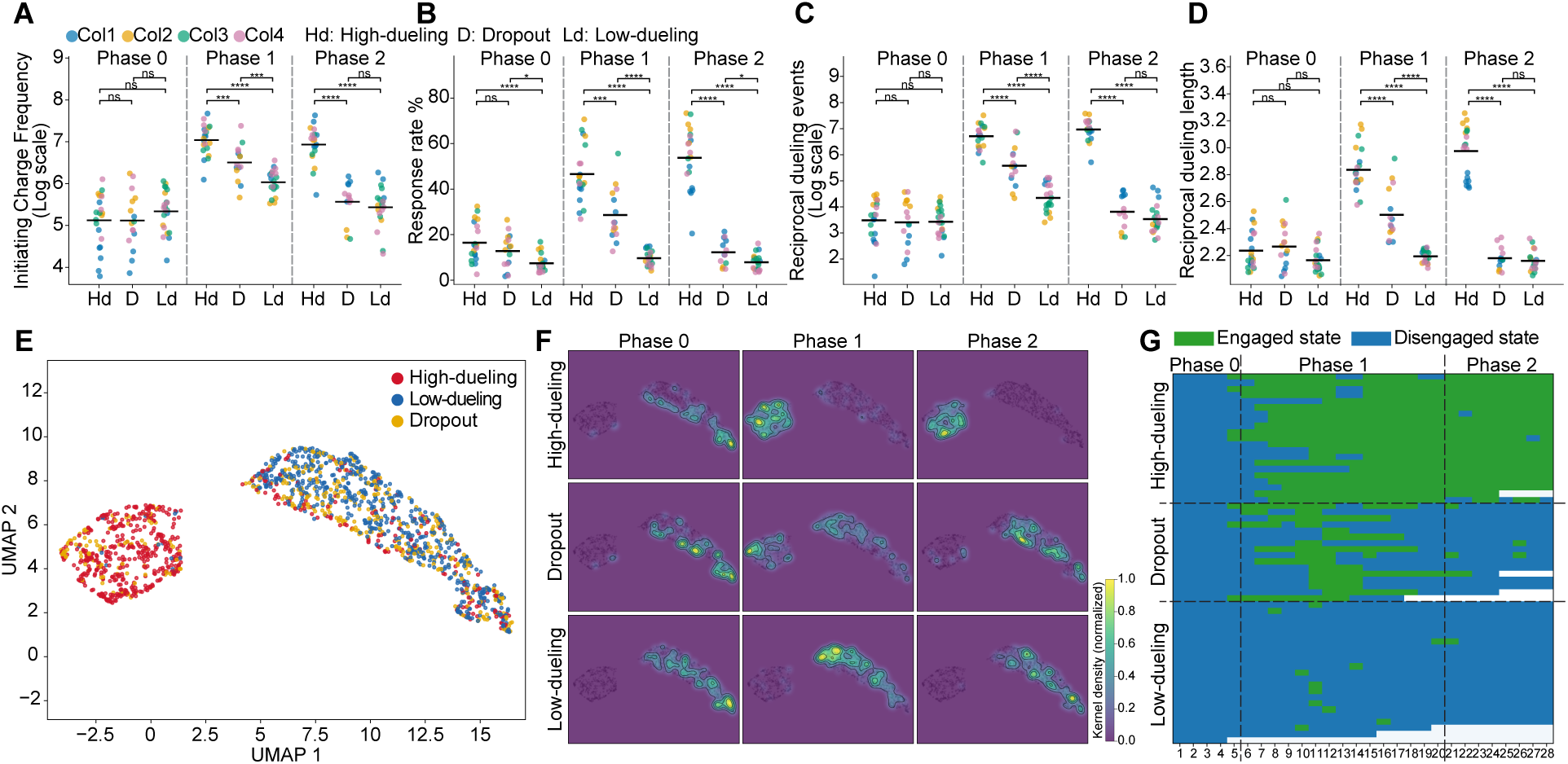
Seven quantitative dueling features reveal two discrete, temporally stable behavioral states. (**A-D**) Phase-stratified dot plots of natural log scaled initiating charge frequency (**A**) and reciprocal dueling events (**C**). Phase-stratified dot plots of response rate (**B**) and reciprocal dueling length (**D**). Each dot represents the phase-averaged charge frequency for one ant, colored by colony (1-4). Horizontal bars indicate colony-pooled means. Significance brackets indicate p values from colony-blocked permutation tests: ns, not significant; **, p < 0.01; ***, p < 0.001; ****, p < 0.0001. (**E**) UMAP embedding of daily seven dueling features. Each point represents one ant on one day, colored by caste-fate trajectory. (**F**) Kernel density maps showing the UMAP-space occupancy of each caste-fate trajectory across the three phases. Each panel is independently normalized. (**G**) Raster plot showing HMM-inferred behavioral-state assignments across days for individual ants from four colonies. Ants within each caste-fate trajectory are ordered as in the heatmap in Figure 1B.

We next assessed whether each initiating charge escalated into reciprocal dueling and quantified duel length as the number of back-and-forth charges exchanged between two ants **(Figure 2C-D)**. Consistent with their elevated response rates, High-dueling ants and Dropouts engaged in more reciprocal dueling than Low-dueling ants in Phase 1, whereas reciprocal dueling occurred predominantly in High-dueling ants during Phase 2 **(Figure 2C)**. Moreover, High-dueling ants exhibited significantly longer duels across Phase 1 and 2, with an average chain length of 3, corresponding to an initiator–receiver–initiator sequence. This indicates that once the target ant charged back, High-dueling ants typically responded with another charge **(Figure 2D)**. These results suggest that responsiveness to received charges reflects a distinct behavioral state associated with gamergate emergence.

### Multidimensional dueling features reveal two temporally stable activity states

Having established an automated tracking pipeline for the antennal-dueling tournament, we next sought to identify latent behavioral states underlying individual dueling trajectories. We extracted seven daily behavioral features from each ant: total charges given, initiating charges given, total charges received, initiating charges received, response rate, reciprocal duel count, and average chain length. We next applied Uniform Manifold Approximation and Projection (UMAP) to visualize the daily behavioral feature space of individual ants (see Materials and Methods; **Figure 2E-F and S2D)**. Intriguingly, the resulting embedding revealed two distinct clusters corresponding to high and low dueling engagement, hereafter referred to as the engaged-state and disengaged-state regions, respectively. High-dueling ants followed a shared trajectory, transitioning from the disengaged-state into the engaged-state region in Phase 1 and remaining there thereafter, indicating that gamergate-fated ants sustain elevated dueling activity throughout the tournament **(Figure 2F)**. By contrast, Dropout ants entered the engaged-state region in Phase 1 but subsequently reverted to the disengaged-state region at the end of Phase 1 and ultimately remained workers **(Figure 2F)**. This pattern indicates that persistence in the engaged-state region, rather than transient entry into it, distinguishes gamergate-fated ants from those that ultimately remain workers.

To test whether the two observed clusters correspond to discrete, temporally persistent behavioral states, we fit a sticky two-state Gaussian hidden Markov model (HMM) ^29^ using the seven dueling features (see Materials and Methods). The model identified two latent states with distinct feature distributions and transition dynamics, with transitions strongly biased toward state persistence, further supporting the existence of temporally stable behavioral states. We designated the state with the higher interaction intensity as the engaged state and the other as the disengaged state **(Figure 2G)**. Overlaying the HMM-inferred states onto the UMAP embedding showed strong concordance between state identity and the two regions in feature space **(Figure S2H)**. In addition, the HMM-defined activity states were highly stable, with 92.9% of adjacent day pairs remaining in the same state (the low-to-low probability = 0.937; the high-to-high probability = 0.914), whereas transitions were infrequent (the low-to-high probability = 0.063; the high-to-low probability = 0.086), indicating that the two clusters correspond to temporally coherent latent states. Consistent with this stability, posterior assignments exhibited near-zero uncertainty (mean entropy = 0.005), indicating highly confident state classification. We further found that once High-dueling ants (future gamergates) transitioned into the engaged state, they stably maintained this state, with only rare and brief 1-2 day shifts back to the disengaged state, likely associated with the feeding schedule. Conversely, Dropout ants entered the engaged state only transiently in Phase 1, then reverted to the disengaged state and remained there throughout Phase 2 **(Figure 2G)**. Together, these analyses indicate that antennal-dueling behavior is organized into two temporally stable activity states and suggest that successful transition to reproductive caste fate requires not only entering but also maintaining the engaged state.

### Engaged-state ants preferentially duel with one another

The Duel-Tracker provided a way to identify the target of each charging event. We next asked whether the three caste-fate trajectories exhibit distinct charge preferences during caste transition. We focused specifically on initiating charges, as these represent an initial decision to engage, rather than response charges that are reactive to a charge. For each ant, we then determined whether its initiating charges were directed toward an engaged-state or disengaged-state opponent (**Figure S3A-B**). Because no ants transitioned into the disengaged state during Phase 0, we backfilled Phase 1 state identities to Phase 0 for the purpose of calculating charge preference. We then calculated a preference index (PI) as the normalized difference between initiating charges directed toward engaged-state and disengaged-state ants, where 0 indicates no preference, 1 indicates charging exclusively toward engaged-state ants, and −1 indicates charging exclusively toward disengaged-state ants. In Phase 0, none of the three caste-fate trajectories showed a clear preference for charging ants in either state (**Figure 3A**). In Phase 1, High-dueling ants and Dropouts exhibited a strong preference for charging engaged-state ants ^21^ (High-dueling ants PI = 0.583 ± 0.019, Dropouts PI = 0.165 ± 0.065), whereas Low-dueling ants exhibited a mild preference for charging disengaged-state ants (PI = −0.344 ± 0.092) (**Figure 3A**). In Phase 2, High-dueling ants retained a strong preference for charging engaged-state ants (PI = 0.706 ± 0.015), whereas Dropouts, like Low-dueling ants, exhibited a mild preference for charging disengaged-state ants (Dropouts PI = −0.222 ± 0.070, Low-dueling ants PI = −0.252 ± 0.061) (**Figure 3A**). Moreover, ants in the engaged state showed a strong preference for targeting other engaged-state ants (**Figure 4C**), indicating that once High-dueling ants and Dropouts transition into the engaged state, they preferentially duel with one another. However, once Dropouts revert to the disengaged state, this preference is lost, and they instead show a mild preference for disengaged-state ants. Next, we quantified the number of initiating charges received by each caste-fate trajectory from engaged- and disengaged-state ants. We found that in Phase 1, both High-dueling ants and Dropouts received significantly more charges from engaged-state ants, whereas in Phase 2, this pattern persisted only in High-dueling ants (**Figure 3B**), further supporting the conclusion that engaged-state ants preferentially duel with one another. In contrast, Low-dueling ants received significantly more charges from disengaged-state ants in both Phases 1 and 2, suggesting that disengaged-state ants also preferentially duel with one another (**Figure 3C**). This strong charging bias, especially among engaged-state ants, suggests that future gamergates may preferentially recognize or interact with one another, forming a positive feedback loop that reinforces engagement during gamergate determination.

**Figure 3.**
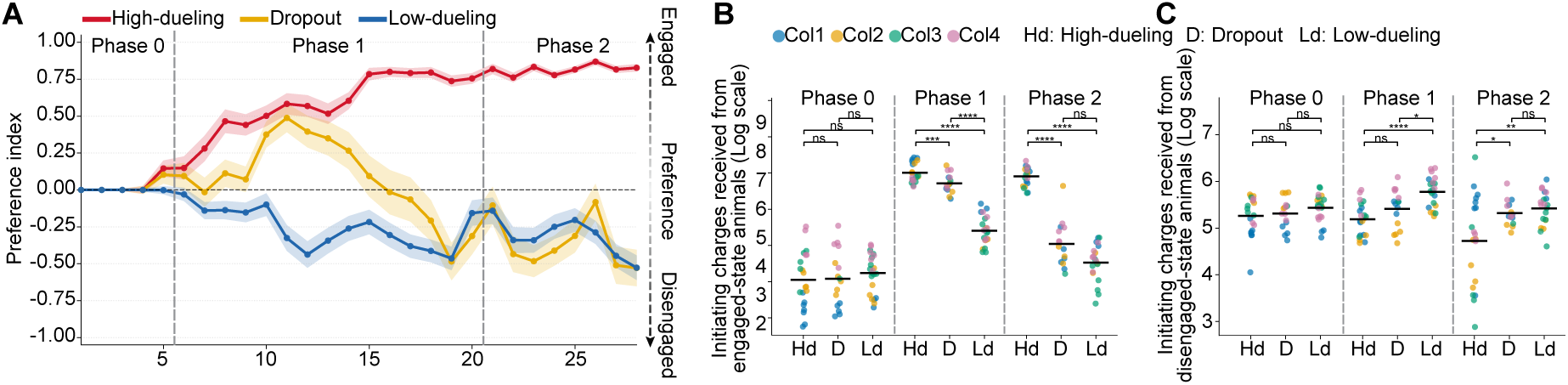
Engaged-state ants show a strong assortative preference by preferentially charging other engaged-state ants. (**A**) Daily preference index (PI) across 28 days for the three caste-fate trajectories (mean ± SEM; four colonies pooled). Dashed horizontal line, PI = 0, no preference. **(B-C)** Phase-stratified dot plots of natural log scaled initiating charges received from engaged-state animals (**B**) or disengaged-state animals (**C**). Each dot represents the phase-averaged daily statistic for one ant. Horizontal bars indicate colony-pooled means. Significance brackets indicate p values from colony-blocked permutation tests: ns, not significant; **, p < 0.01; ***, p < 0.001; ****, p < 0.0001

**Figure 4.**
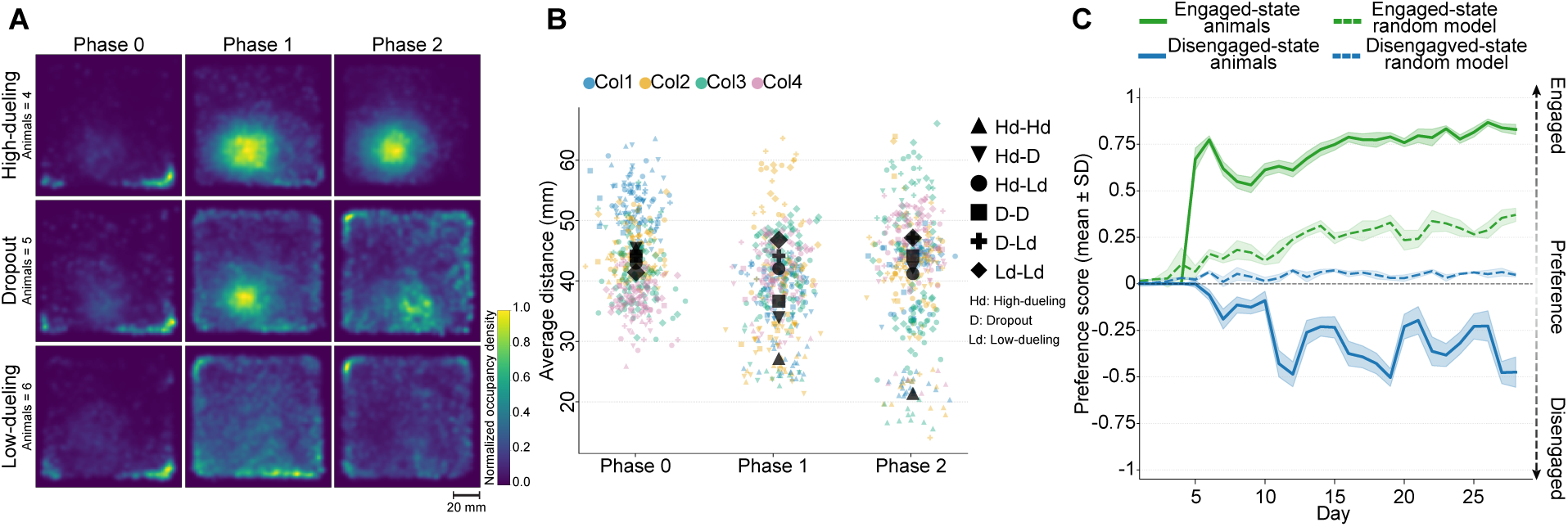
Engaged-state ants form a spatially segregated dueling circle, yet their assortative preference exceeds predictions from random-collision models based on spatial proximity alone. (**A**) Kernel density maps showing arena-space occupancy for the three caste-fate trajectories across phases in Colony 4. Scale bar = 20 mm. (**B**) Phase-stratified dot plots of pairwise Euclidean distances between ants from the three caste-fate trajectories. Each dot represents one pair of ants from the same colony. Black markers indicate colony-pooled means. **(C)** Daily preference index (PI) across 28 days for engaged- and disengaged-state ants (mean ± SEM; four colonies pooled). Dashed lines indicate the expected PI from spatially constrained random-collision null models.

### A spatially segregated dueling circle emerges during caste transition

We observed that engaged-state ants showed a strong assortative preference for dueling with one another once antennal dueling intensified in transition colonies, raising the possibility that engaged-state ants recognize one another. An alternative explanation is simple co-aggregation, whereby engaged-state ants occupy the same region of the arena and thus interact more frequently. To test this possibility, we quantified spatial occupancy across the arena for the three caste-fate trajectories across the three phases **(Figure 4A and S4A-C)**. In Phase 0, all ants either aggregated in one corner of the arena or were diffusely distributed, reflecting nest exploration and a lack of clear social preferences. Strikingly, in Phase 1, High-dueling ants and Dropouts began to aggregate at a single location, forming a “dueling circle” described in previous studies ^16,30^. However, in Phase 2, Dropouts returned to the disengaged state and left the dueling circle, occupying the periphery like Low-dueling ants, indicating that the strong assortative preference for dueling among engaged-state ants cannot be explained by simple co-aggregation alone **(Figure 4A and S4A-C)**. In addition, we measured average pairwise distances among the three caste-fate trajectories across the three phases **(Figure 4B)**. Pairwise distances were indistinguishable among ants in Phase 0. Consistent with the occupancy results, in Phase 1 High-dueling ants were closest to one another (26.79 ± 1.44 mm), followed by High-dueling ants with Dropouts (32.12 ± 2.65 mm) and Dropouts with Dropouts (36.63 ± 2.80 mm). The substantially greater distances between Low-dueling ants and other ants (43.28 ± 0.79 mm) indicate their peripheral localization and spatial separation from each other and from the dueling circle **(Figure 4B).** Together, the state-specific interaction preferences and spatial segregation provide additional evidence that the engaged and disengaged states reflect biologically meaningful behavioral states and may correspond to distinct internal states **(Figure S4D)**.

Engaged-state ants sharply increased preferential charging toward one another in Phase 1 **(Figure 4C)**, raising two possibilities: *(i)* this pattern arises solely from spatial proximity, or *(ii)* engaged-state ants exhibit mutual recognition that biases interactions beyond spatial proximity alone. To distinguish between these possibilities, we implemented a spatially constrained random-collision model, conceptually similar to spatial null models used to disentangle social preference from encounter structure (Richardson et al., 2015), in which the probability of interaction between two ants was assumed to be inversely proportional to their pairwise distance, thereby predicting the charging preference expected from spatial proximity alone. We found that the observed preference significantly deviated from the random-collision model prediction in both states, being higher than predicted for engaged-state individuals and lower than predicted for disengaged-state individuals (**Figure 4C**; P < 0.0001), suggesting that state-dependent interaction patterns cannot be explained by spatial proximity alone and may involve mutual attraction or recognition mediated by chemosensory cues.

### Early access to abundant food biases antennal dueling and promotes gamergate fate

In colonies composed of age-matched workers from the same source, gamergate-fated ants rapidly shifted into engaged state and preferentially targeted other engaged-state ants in Phase 1. This early assortative targeting suggests that differences in behavioral or physiological state before transition starts may precede overt caste divergence and may bias competitive success during caste transition. It appears that nutritional state is a compelling candidate: antennal dueling is energetically costly, and dueling intensity follows feeding cycles in our data **(Figure S5A)**. Mechanistically, genes in nutrient-sensing pathways, particularly insulin/IGF signaling, are differentially expressed and are higher in gamergates vs. workers in *H. saltator* and insulin administration promotes ovarian development without increasing dueling behavior ^23^, suggesting a route through which diet may shape fat-body metabolism, vitellogenin production, and possibly the behavioral persistence required to prevail during caste transition. To test how nutritional state affects caste transition, we established transition colonies under two feeding regimes using callows from the same source colony: ad libitum feeding (every 2 days) and a calorie-restricted feeding regime (every 5 days, as in previously described experiments). On day 6, after both groups had been fed the previous day, we mixed similar numbers of ants from the two groups, tracked their behavior, and thereafter maintained all mixed colonies on the same feeding schedule (every 5 days). Strikingly, 18 of 23 of previously ad libitum-fed ants became gamergates, compared with 7 of 24 previously calorie-restricted ants across four independent mixed transition colonies **(Figure 5)**. Consistent with this, quantification of dueling activity revealed that more ad libitum-fed ants shifted into and remained in the engaged state than calorie-restricted ants (p = 0.015) **(Figure 5A-B)**. In addition, engaged-state ants in both feeding regimes showed a strong charging preference toward other engaged-state ants, whereas disengaged-state ants showed no clear preference (**Figure 5C**). This result suggests that nutrient status primarily biases entering the engaged state, rather than altering assortative dueling preferences. Notably, engaged-state ants in the calorie-restricted group exhibited greater variation in PI than those in the ad libitum group (p=0.013). Moreover, head width did not differ significantly between feeding groups **(Figure S6E)**, indicating that these effects are not explained by body size differences, which is established during larval life as adults do not grow. Together, our results indicate that previous access to abundant food shapes dueling activity and reinforces dominance, conferring an advantage in the transition to gamergate fate.

**Figure 5.**
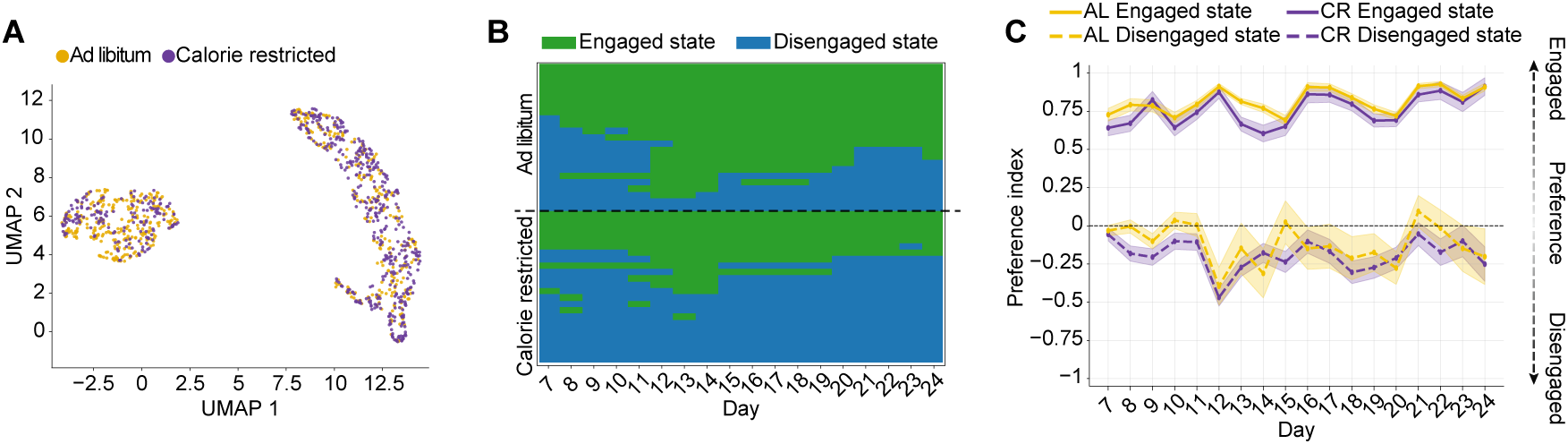
Early increased nutrition biases entry into and persistence of the engaged state and transition into gamergates. (**A**) UMAP embedding of daily seven dueling features. Each point represents one ant on one day and is colored by feeding regime. (**B**) Raster plot showing HMM-inferred behavioral-state assignments across days for individual ants from the two feeding regimes. (**C**) Daily preference index (PI) after ants from the two feeding groups were mixed (mean ± SEM; four colonies pooled). Solid line represents engaged-state ants. Dashed line represents disengaged-state ants.

### Aggressive biting shapes social dominance and caste determination

The automated tracking pipeline provided unprecedented resolution of individual dueling dynamics and revealed that Dropouts, which represented approximately one-fourth of ants participating in the tournament, engaged intensely in dueling during Phase 1 but subsequently withdrew and remained as workers. When Dropouts entered the engaged state in Phase 1, their daily dueling features were indistinguishable from those of High-dueling ants that later became gamergates (silhouette score = 0.0503; **Figure 2E**). What factors constrain their caste fate? Previous studies have shown that aggressive behaviors, such as policing bites, can trigger reversion of gamergates to workers when an external gamergate is introduced in a colony that has an established reproductive caste ^15,17^. During the transition, we frequently observed two distinct biting behaviors: aggressive grooming and mandible locking. *(i)* Aggressive grooming consisted of repeated use of the maxillolabial complex to groom the opponent, typically followed by biting the head while dragging the target toward the initiator **(<u>Video S4</u>)**. This interaction exhibited clear dominant and submissive roles. Intriguingly, we frequently observed the submissive ant becoming immobile in a freezing-like response after being aggressively groomed (**<u>Video S4</u>; Figure S6A**). *(ii)* Mandible locking is a prolonged reciprocal grapple in which two ants clamp onto one another with their mandibles and wrestle for extended periods, sometimes exceeding one hour **(<u>Video S5</u>)**. Dominance was assigned to the initiator as the dominant individual (biter) and the other ant as its target during mandible locking as the target often resisted and maintained the hold. To systematically quantify these biting behaviors, we trained a neural-network classifier to detect aggressive grooming and mandible locking by learning spatiotemporal features from manually annotated events (see Materials and Methods). We then manually curated the detected events to quantify event frequency and bout duration. Aggressive grooming occurred much more frequently than mandible locking (448.8 ± 179.6 *vs.* 45.5 ± 11.7 events per colony) (**Figure 6A-B**). In addition, aggressive grooming exhibited a shorter range of bout durations, spanning 10 s to 10 min, compared with 10 s to 30 min for mandible locking (**Figure S6A-B**). Intriguingly, these biting behaviors were concentrated in Phase 1, when High-dueling ants and Dropouts entered the engaged state with intense dueling activity. Aggressive grooming was directed toward both Dropouts and Low-dueling ants (37.7% versus 56.6%), whereas mandible locking was more often directed toward Dropouts (44.9%). In contrast, High-dueling ants (future gamergates) received very few biting events (5.7% for aggressive grooming, 7.6% for mandible locking). Body size appeared to play only a limited role in aggressive grooming. Initiators had larger head widths than their targets in 51.8% of events and were on average only 0.42% larger (Figure S6C). By contrast, mandible-locking initiators were larger than their targets in 71.8% of events and were, on average, 2.51% larger (**Figure S6D**). To determine how these two biting behaviors contribute to social hierarchy in transition colonies, we calculated Elo scores ^31^ for each ant throughout the caste transition. Elo scores iteratively update an individual’s dominance estimate based on the outcomes of pairwise interactions, such that individuals gain points when winning (initiator) and lose points when defeated (target), allowing dynamic tracking of relative social rank over time. Elo trajectories derived from aggressive grooming showed that High-dueling ants and Dropouts occupied the highest rank percentiles from early Phase 0 to approximately day 15, after which Dropouts declined sharply and stabilized at intermediate ranks between High-dueling ants and Low-dueling ants (**Figure 6C**). In contrast, Low-dueling ants rapidly fell to low rank percentiles in Phase 0 and remained there throughout the caste transition (**Figure 6C**). Elo trajectories derived from mandible locking revealed a distinct pattern: the three caste-fate trajectories were largely indistinguishable in Phase 0 because few events occurred but gradually diverged during Phase 1. After day 15, Dropouts showed a sharp decline in rank percentiles and eventually fell below Low-dueling ants due to receiving more mandible locking interactions (**Figure 6D**).

**Figure 6.**
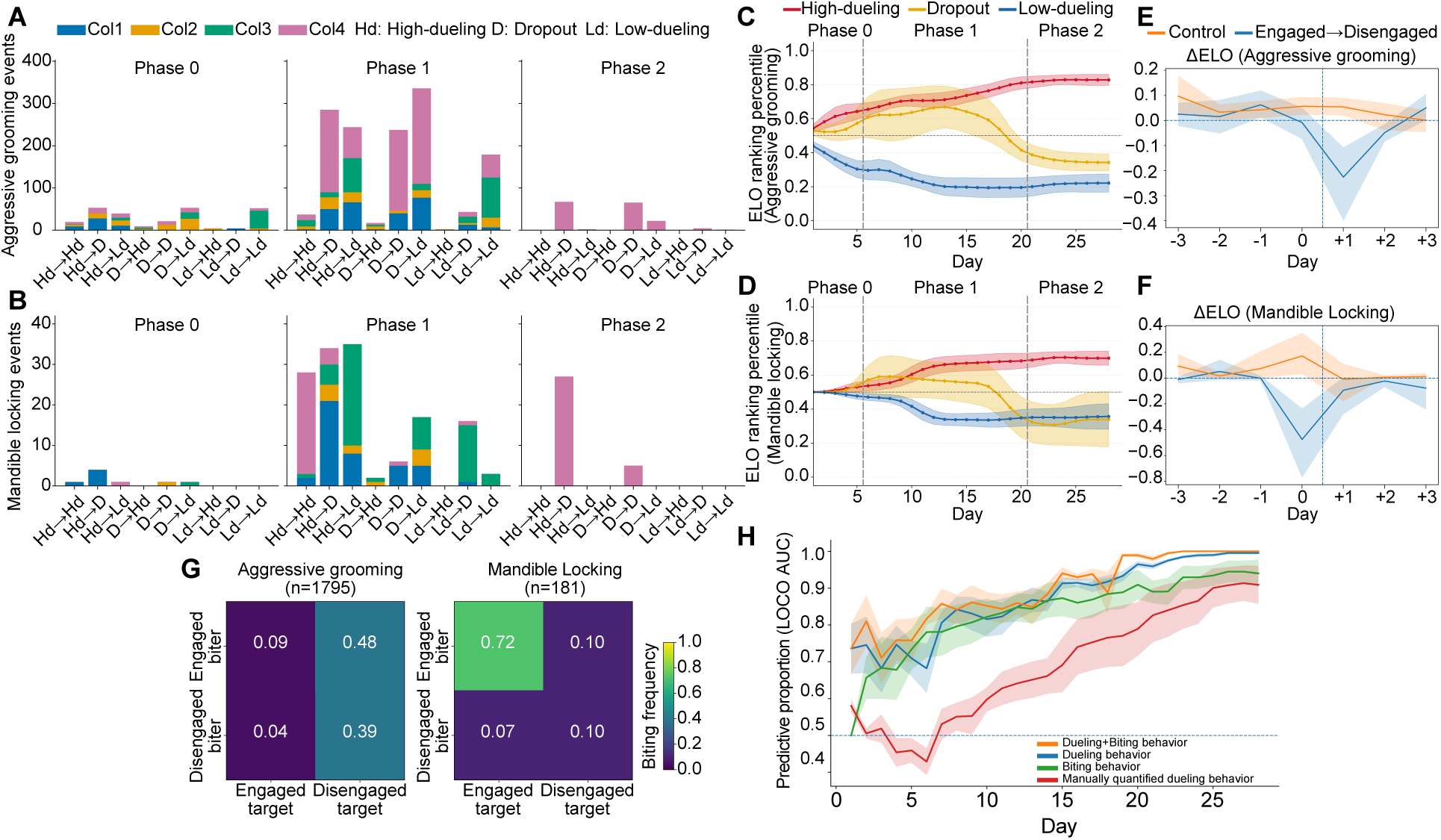
Aggressive grooming and mandible locking shape dominance rank and are associated with withdrawal from the reproductive trajectory. (**A-B**) Stacked bar histograms showing counts of aggressive grooming (**A**) and mandible locking (**B**) across all interaction directions between pairs of caste-fate trajectories. (**C-D**) Daily Elo rank percentiles for the three caste-fate trajectories across 28 days, derived from aggressive grooming (**C**) and mandible locking (**D**) interactions (mean ± SEM). (**E-F**) Peri-transition analysis of day-to-day changes in Elo rank percentiles derived from aggressive grooming (**E**) and mandible locking (**F**) interactions (ΔElo; mean ± SEM), aligned to the engaged-state day (day 0) before transition to the low state (day +1). (**G**) Pairwise interaction frequencies for aggressive grooming and mandible locking between engaged- and disengaged-state ants, pooled across four colonies. (**H**) Predictive performance of elastic net-regularized logistic regression models for predicting caste fate (gamergate vs. worker) across recording days (leave-one-colony-out AUC; mean ± SEM; n = 4 colonies).

The sharp decline in both Elo ranks in Dropouts closely coincided with their withdrawal from intense dueling. To examine how these two biting behaviors relate to this shift, we focused on ants transitioning from the engaged state to the disengaged state and quantified changes in their Elo scores within a ±3-day window. Intriguingly, we observed that Elo derived from aggressive grooming decreased only after the transition, whereas Elo derived from mandible locking decreased prior to state transition (**Figure 6E-F**). This temporal dissociation suggests a sequential mechanism in which mandible locking precedes the shift from the engaged state and contributes to destabilizing it, whereas aggressive grooming follows the transition and reinforces the disengaged state. In addition, aggressive grooming was directed predominantly toward disengaged-state individuals, which comprised 87% of targets, and biters included both engaged-and disengaged-state ants (48% and 39%, respectively). In contrast, mandible locking was highly concentrated between engaged-state ants (72%) (**Figure 6G**), consistent with ranking percentiles declining prior to state transition. Next, we trained elastic net-regularized logistic regression models to evaluate the predictive power of dueling and biting behaviors (see Materials and Methods; **Figure 6H**). Models based on automated tracking features outperformed the previously described manual quantification proxy ^21^. Notably, the model incorporating both dueling and biting features achieved the highest predictive performance (AUC = 0.95 on day 20). Together, these analyses reveal that reproductive hierarchy formation is not explained by dueling frequency alone. Instead, caste fate emerges from the stability of a behavioral state embedded within a broader social regulatory system: reciprocal antennal dueling sustains engagement among future gamergates, mandible locking precedes withdrawal from this state, and aggressive grooming reinforces the resulting worker trajectory. Thus, multiple interaction modes act at distinct phases of the tournament to shape which workers persist toward gamergate fate.

## Discussion

Caste fate in most eusocial insects is developmentally fixed ^32^, limiting opportunities to examine hierarchy establishment in adults. We studied adult hierarchy formation in *H. saltator* by analyzing age-matched workers undergoing transition to reproductive gamergates in queenless colonies. Because gamergates and workers are genetically and morphologically similar, differing primarily in physiology and behavior, hierarchy is established and maintained through social interactions coupled with internal physiological signals and neural mechanisms. Here, we used the NAPS automated tracking system ^26^, together with machine learning classifiers, to quantify these interactions and test how they shape social rank. We quantified three social behaviors: antennal dueling, aggressive grooming, and mandible locking, each with a distinct role in establishing the reproductive hierarchy during colony transition.

### Winner-winner effect in ritualized antennal dueling

Ritualized antennal dueling is closely associated with caste transition in *H. saltator* colonies and can persist for more than one month ^19^. However, its function has long been debated. Prevailing theories propose that dueling either acts as a “winner-winner” interaction that progressively reinforces reproductive status ^33^, or reflects a tournament with an early winner-loser phase, in which most workers engage dueling but rapidly withdraw, followed by a later winner-winner phase in which dueling becomes concentrated among future gamergates ^16^. Here, we used an automated tracking pipeline to decompose dueling into spontaneity, attractiveness, responsiveness, and reciprocity, and identified two attractor-like behavioral states underlying these dynamics (**Figure 2A**). We found that once an ant shifted into the engaged state, it preferentially targeted other engaged-state ants and tended to remain in that state over time. Moreover, High-dueling ants and Dropouts exhibited indistinguishable daily dueling features while in the engaged state, supporting the idea that winner-winner interactions among engaged-state ants reinforce dominance. Prior observations show that dopamine levels are elevated in gamergates relative to workers in *H. saltator*, and that dopamine declines rapidly in workers policed by nestmates during the caste transition ^21^. The intense dueling we observed, characterized by higher spontaneity and responsiveness, may reflect elevated neuromodulatory activity involving dopamine and octopamine signaling, which regulate arousal and social responsiveness in *Drosophila* ^34,35^. Frequent reciprocal dueling in the engaged state likely forms a positive feedback loop that stabilizes state maintenance. This winner-keep-winning phenomenon has also been observed in male mice, where repeated victories progressively enhance aggressiveness through plasticity in the VMHvl aggression circuit ^36^, and dopamine further modulates aggression in an experience-dependent manner ^37^. More broadly, reciprocal social interactions can alter endocrine state and reinforce behavioral engagement across taxa, including dopamine-associated reproductive dominance in bumble bees ^38^, juvenile hormone-associated dominance in paper wasps ^39^, and androgen-mediated winner effects in cichlid fish ^40^.

### Mutual recognition and spatial segregation

In *H. saltator*, active dueling ants have been described as forming a “gamergate circle” during caste transition, with dueling concentrated in a fixed region of the nest ^30^. This pattern is consistent with our finding that ants in the engaged state form a dueling circle, while Dropouts subsequently transition to the disengaged state and leave this region. In parallel with this aggregation of engaged-state ants, disengaged-state ants tend to avoid the dueling circle (**Figure 4B**), leading to rapid spatial segregation at phase 1. Such spatial structuring suggests that state-dependent interactions actively organize the social space within the colony. Spatial segregation of reproductively active individuals is a common feature in social insects, where nests often contain distinct regions specialized for brood rearing, food storage, and reproductives ^32,41^. This raises the possibility that the dueling circle corresponds to a central reproductive zone in which prospective gamergates preferentially localize. These patterns suggest that mutual recognition among prospective reproductives drives preferential interactions among engaged-state individuals and leads to their spatial segregation, consistent with observations in other ant species such as *Lasius niger*, where fertility-associated chemical signals correlate with reproductive status and influence the outcome of reproductive conflicts ^42^. In engaged-state ants, such recognition is likely mediated by olfactory cues, as visual recognition is limited in the dark conditions of natural nest environments ^17^. This dynamic, state-dependent recognition likely corresponds to changes in cuticular hydrocarbon (CHC) profiles during the caste transition. Gamergates show increased expression of lipid metabolism and modifying enzymes, along with a shift in cuticular hydrocarbon profiles toward longer-chain hydrocarbons ^17,23^. Detection of these chemical cues is mediated through antennal olfactory pathways and is required for normal social interactions, as disruption of olfactory signaling (e.g., via Orco mutagenesis) impairs social behavior ^43,44^. Furthermore, antennae harbor both olfactory and mechanosensory neurons, and studies in moths show that mechanical stimulation can signal the release of neuropeptides that regulate pheromone biosynthesis ^45^. Antennal dueling may act as sensory stimulation that shifts cuticular hydrocarbon profiles, thereby signaling reproductive status ^17^.

### Nutritional state modulates antennal dueling

In social insects, diet plays a pivotal role in determining caste fate and behavior. For example, in honeybees (*Apis mellifera*), larvae destined to become queens are fed a nutrition-rich royal jelly that induces queen development via epigenetic mechanisms involving DNA methylation ^46^. Similarly, in the harvester ant (*Pogonomyrmex rugosus*), the presence of trophic eggs has been shown to induce queen development ^47^. Beyond developmental effects, diet shapes social behavior in adult insects. In honeybees, a diet rich in protein (pollen) increased levels of vitellogenin, which modulates behavior and acts as an antioxidant, while reducing expression of stress proteins like HSP70 ^48,49^. Additionally, pollen intake increases brain amino acid levels over time, potentially supporting neural function and behavior by influencing neurotransmission and neural plasticity ^50^. In Argentine ants (*Linepithema humile*), depriving colonies of carbohydrates significantly reduces aggression and activity, primarily by diminishing worker fat stores ^51^. In *H. saltator*, gamergates exhibit elevated brain-derived insulin signaling that activates the MAPK pathway in the fat body, increasing lipid and vitellogenin production and inducing development of the ovary. The reactivated ovary then secretes an insulin antagonist, Imp-L2, that inhibits the aging-associated AKT branch, thus allowing reproduction without compromising longevity ^23^. Here, we show that early access to abundant food confers a strong advantage in entering and maintaining the engaged state and in attaining gamergate status (**Figure 5**). Moreover, antennal dueling intensified immediately after the first feeding, suggesting nutritional state may bias neural circuits toward dueling. However, insulin alone is not sufficient to induce antennal dueling, as insulin injection promoted ovariole development but did not increase dueling ^23^.

### Hierarchy formation through distinct types of dominance behaviors

Aggressive behavior is a widespread mechanism for establishing dominance across both vertebrates and invertebrates ^52^. In social insects, policing is a common strategy that suppresses worker reproduction, often through aggression toward egg-laying workers and/or removal of worker-laid eggs, thereby preserving reproductive hierarchy and division of labor ^53,54^. Close-contact interactions involving antennation, mandible contact, and sustained physical manipulation have been described in ants and wasps as components of dominance interactions and worker policing ^41,44^. We systematically quantified two forms of dominance biting, aggressive grooming and mandible locking, and uncovered their distinct roles during caste transition in *H. saltator*. Aggressive grooming showed clear asymmetry and was associated with rapid hierarchy formation in Phase 0 (**Figure 6C**), consistent with the dominance-based hierarchy formation reported in another Ponerine ant, Dinoponera ^55^ and paper wasps ^56^. We speculate that this behavior stabilizes the disengaged state through neuromodulatory and endocrine pathways, potentially involving reduced dopaminergic and juvenile hormone signaling, together with CHC-mediated social feedback, thereby creating a reinforcing loop that maintains subordinate behavioral states ^57–59^. We frequently observed that ants receiving aggressive grooming entered a transient freezing-like state (54.1%). This behavior is reminiscent of submissive crouching observed in ant dominance interactions and death feigning reported in ants during interspecific conflicts ^60^. More broadly, it is consistent with vertebrate social-defeat paradigms, in which freezing and immobility often accompany passive subordinate coping ^61^.

Mandible locking occurs much less frequently than aggressive grooming and is concentrated in Phase 1, when intense dueling occurs. Most mandible locking events occurred between engaged-state ants (**Figure 6G**), and intense reciprocal dueling occasionally escalated into mandible locking, suggesting that this behavior serves as a tie-breaking mechanism when two engaged-state individuals occupy similar hierarchical positions. Initiators of mandible locking also tend to be larger, consistent with broad findings in ants that larger individuals often have a higher probability of winning aggressive interactions and attaining higher dominance rank ^32^. In addition, we observed that Dropouts often transition from the engaged state to the disengaged state after receiving mandible locking, followed by aggressive grooming. We speculate that mandible locking may involve deposition of a chemical mark on Dropouts, similar to the pretender punishment reported in the queenless ant *Dinoponera quadriceps*, where the alpha female uses Dufour’s gland secretion to mark challengers and elicit worker aggression ^15^.

### Antennal dueling as a potential honest signal of reproductive potential

Animals frequently display physical condition to secure reproductive opportunities. A classic example is the peacock’s train: only high-condition individuals can sustain energetically costly ornaments, making them reliable signals of reproductive fitness ^62,63^. We propose that antennal dueling may function as an honest signal linking individual fitness to reproductive opportunity during caste transition. Dueling acts as a dynamic behavioral broadcast maintained under sustained metabolic and social pressure. Only a subset of workers maintains the engaged state and ultimately become gamergates, whereas others withdraw or fail to persist. In social insects, fertility is often communicated through cuticular hydrocarbons (CHCs) that correlate with ovarian activity ^17^. However, chemical signals can be vulnerable to cheating or mimicry ^64,65^. In contrast, behavioral signals require continuous performance and thus impose ongoing energetic costs. Notably, antennal dueling persists for weeks, whereas ovary activation occurs within approximately 10 days, suggesting that prolonged interaction functions as a robustness filter rather than simply driving reproductive development. From an evolutionary perspective, antennal dueling represents a form of ritualized aggression that enables mutual assessment while minimizing injury. Across taxa, ritualized contests such as roaring in red deer allow repeated evaluation without immediate escalation ^66,67^, although conflicts may intensify when individuals are closely matched. Similarly, dueling in *H. saltator* can escalate into mandible locking, indicating a graded conflict system. Compared to species that rely on injurious aggression to establish hierarchy ^55^, intensive dueling is restricted to a smaller group and allows withdrawal, reducing unnecessary energy expenditure and worker loss. Together, antennal dueling constitutes a temporally extended, costly, and socially reinforced honest signal of fecundity that potentially promotes reliable selection of reproductively competent individuals, thereby promoting colony-level reproductive efficiency.

## Acknowledgments

We are grateful to Rory Coleman for comments and suggestions on the manuscript. We thank Scott Wolf, Ian Traniello, Dee Ruttenberg from the Kocher lab for providing valuable insights of setting up NAPS. We thank Sarah Kocher and Hua Yan for comments and suggestions on the project. We thank Darren Lin and Cheryl Kim for annotating biting behaviors. We thank Cecilia J Reisner for the technical assistance in the laboratory and all the support from the Desplan lab.

## Materials and Methods

### Animal rearing and tagging

We performed four independent replicates, including both natural-transition and nutrition-manipulation experiments using the ant *Harpegnathos saltator*. Each colony consisted of 15 callow workers from the same source colony housed within a 98 x 98 mm acrylic arena lined with a ∼20 mm layer of plaster to maintain humidity. Colonies were maintained inside a 25 °C recording tent under a 12 h light: 12 h dark photoperiod, illuminated by LED light bars to mimic natural circadian rhythms. All experiments were conducted between November 2024 and October 2025. Individual identification was achieved using ArUco tags from the 3×3_50 library (Garrido-Jurado et al., 2014), printed on waterproof, tear-resistant paper (Revolution NeverTear) and cut to 2.5 x 2.5 mm. Prior to tagging, callows were briefly anesthetized on wet ice in an acrylic box until movement ceased. Each tag was hand-cut and affixed to the thorax using UV-curable resin glue (Let’s Resin). When properly applied, the resin glue did not affect ant mortality or behavior. Following recovery, tagged ants were returned to the arena, where colonies were provided food every five days in the form of three medium-sized crickets.

### Recording and Tracking

Arenas were constructed from clear acrylic boxes (98 mm x 98 mm) with a thin layer of plaster at the base to maintain humidity. Colonies were recorded beginning at least 12 hours after establishment to ensure behavioral stabilization. Video acquisition was performed from above using a Basler a2A1920-160ucBAS camera capturing 2000 px x 2000 px frames at 30 frames per second, 12 hours a day from 7 am to 7 pm. Recordings were compressed in real time using CAMPY, a Python-based package for high-efficiency video acquisition and compression. The resulting videos had a spatial resolution of approximately 0.05 mm per pixel, allowing high-fidelity tracking of fine-scale interactions. Pose estimation was performed using SLEAP ^68^ with a 17-node skeleton marking the head, thorax, tag, abdomen, abdomen tip, left and right mandibles, left and right antennal joints, and three pairs of legs. The model was trained on 200 manually labeled frames, achieving 95% of node localization errors within 8.5 pixels in the validation set. Individual identities were subsequently resolved using NAPS ^26^, which integrates SLEAP pose tracking with ArUco tag recognition to maintain consistent individual labeling throughout recordings.

### Post-processing and Filtering

After pose estimation and identity assignment, a multi-stage filter was applied to remove implausible detections while retaining natural rapid movements characteristic of dueling behavior. (1) Node exclusion: six leg nodes were discarded from all analyses due to frequent occlusion and inconsistent tracking. (2) Unused tags: detections mapped to tag IDs not used in the experiment were removed. (3) Velocity outliers with a coordinated-motion safeguard: per-node velocities were computed from frame-wise position gradients. A node was flagged as a velocity outlier if its instantaneous speed exceeded 10 pixels per frame; however, frames with coordinated motion (>= 2 nodes of the same individual simultaneously exceeding the threshold) were retained to avoid filtering true body movements. (4) Antenna-tip relaxation: to preserve rapid antennal sweeps and contact events, antenna-tip nodes were filtered with a more permissive threshold (up to 200 pixels per frame) relative to other body parts. (5) Skeleton-edge irregularities: for every pair of nodes within an individual, inter-node distances were computed across time; frames with a z-score > 5 relative to that edge’s distribution were treated as geometric outliers, and both nodes forming the affected edge were removed from those frames. (6) Spatial overlaps across individuals: to prevent duplicated assignments, if any node from two different individuals was within 1 pixel in a frame, both nodes were set to missing at that time point. Finally, adaptive interpolation was performed: gaps up to 10 frames were linearly interpolated, and any remaining longer or edge gaps were filled by the nearest available value, ensuring continuous but artifact-free trajectories for downstream behavioral analyses.

### Duel-tracker

To identify dueling events from continuous behavioral recordings, a spatiotemporal graph convolutional neural network (ST-GCN) classifier ^28^, designated Duel-Tracker v3, was developed. Dueling was decomposed into single charging events, in which an ant moved toward a target while rapidly striking with its antennae. Pose data from SLEAP-tracked videos were converted into 60-frame (2 s) overlapping windows sampled every 15 frames (0.5 s), each containing the ego individual and its nearest target. The nearest target for each ego was determined dynamically by a voting algorithm that combined the positions of the mandibles to identify the body node of other individuals most frequently in closest proximity. Each window was then realigned to the ego’s body axis and normalized by body length and orientation. For model training, dueling timestamps were manually annotated within the first 10 min of 13 videos spanning distinct timepoints and feeding contexts, resulting in more than 2,000 dueling events. To improve training robustness, a larger proportion of training clips were sampled from periods immediately following feeding, where ants exhibit increased aggregation and node overlap, posing challenges for pose-based interaction detection. In contrast, the held-out test set was distributed more uniformly across time, providing a more realistic evaluation of model performance under typical behavioral conditions. Rolling windows centered on annotated events were treated as positive samples, whereas non-overlapping windows from unlabeled intervals were used as negative samples. Twenty percent of the labeled videos were withheld as a held-out test set. To mitigate class imbalance, negative windows were down-sampled to match the number of positive windows, and training used label-smoothed binary cross-entropy loss. The ST-GCN operated on an ego–target pose graph containing 11 ego joints and one target node. Models were trained on a GPU with early stopping. For inference, overlapping positive windows were clustered into event-level detections to generate time-resolved duel predictions for each individual.

The final model achieved an ROC-AUC of 0.997 on the training set and 0.93 on the held-out test set, with window-level F1 scores exceeding 0.73 for training and 0.80 for testing after probability-threshold screening. Original script and sample data can be found at: https://github.com/sz-George-l/naps-duel-tracker.

### Reciprocal dueling analysis

Reciprocity between individual pairs was quantified based on direction-specific duel initiation. Each detected duel was represented as a directed event from an initiator (A) to a target (B). Empirical analysis of inter-duel intervals showed that over 95% of responses occurring within 30 s took place within 2 s; therefore, a response charge was defined as a target-specific counter-duel (B -> A) occurring within 2 s of the initial event (A -> B). Duels that did not receive such a response were classified as initiating charges. For events that elicited responses, temporally contiguous, alternating interactions between the same pair were grouped into reciprocal chains (e.g., A -> B -> A -> B …). Chain length was defined as the total number of events in the sequence. Each reciprocal chain was treated as a single reciprocal interaction event, and both participants were assigned one reciprocal event count per chain, regardless of chain length.

### UMAP and HMM analysis

To visualize daily behavioral variation, a 2D embedding was generated using Uniform Manifold Approximation and Projection (UMAP; McInnes et al., 2018). For each individual-day, seven dueling-related features were extracted: hourly average charges given, hourly average charges received, response rate, mean reciprocal chain length, reciprocal duel count, initiating charge count, and received-initiative count. Before dimensionality reduction, features were z-score standardized. UMAP was performed with Euclidean distance, 30 nearest neighbors, a minimum distance of 0.15, and a fixed random seed (random_state = 42). To validate whether the clustering structure reflected discrete behavioral modes, a sticky two-state Gaussian hidden Markov model (HMM) was fitted using hmmlearn ^29^. Each individual-day was assigned to one of two latent behavioral states while temporal dependence between consecutive days was incorporated. The transition matrix was initialized with a self-transition probability of 0.85. The state with the higher combined mean of hourly charges given and received was designated as the engaged state, whereas the lower-interaction state was designated as the disengaged state.

### Phase boundaries and Trajectory assignment

The experimental timeline was divided into three phases relative to colony establishment: Phase 0 (days 1-5), Phase 1 (days 6-20), and Phase 2 (days 21-28). These phase boundaries were chosen to capture the early low-activity period, the expansion phase of dueling engagement, and the late-stage stabilized period. Each individual was assigned a trajectory label based on its pattern of engaged state occupancy across phases. Individuals that entered the engaged state during Phase 1 and maintained occupancy into Phase 2 were classified as High-dueling trajectory. Individuals that occupied the engaged state during Phase 1 but exited this state by the end of Phase 1 were classified as Dropouts. Individuals that did not exhibit sustained engaged-state occupancy were classified as Low-dueling trajectories.

### Preference score and spatially constrained null model

A daily preference score was computed for each individual using the formula: 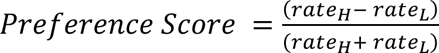, where rate_H and rate_L denote interaction rates toward engaged- and disengaged-state individuals. Rates were calculated as the number of interactions directed toward partners in each state divided by the number of available partners in that state. To determine if observed preferences were explained by spatial proximity, a random-collision null model was constructed where interaction propensity (w_ij) was defined as 1 / d_ij ^69^. The total expected interaction propensity toward engaged- and disengaged-state partners was then computed and normalized by partner abundance to calculate the expected (null) preference score for direct comparison with observed data.

### Caste fate assignment

To assign caste fate, we integrated physiological and molecular measurements, including yolky oocyte number and the expression levels of vitellogenin (Vg) and corazonin (Cz). At day 28, individuals were frozen and dissected, and ovaries were removed to quantify the number of yolky oocytes (yolk-filled structures). In parallel, Vg and Cz transcript levels were measured from brain tissue using quantitative PCR (qPCR) and normalized to internal housekeeping genes. Individuals were clustered using k-means (k = 2) based on these features. The cluster with higher average yolky oocyte number was designated as the gamergate caste, consistent with established markers of reproductive status ^22,23^. This classification showed strong concordance with behavioral trajectories: 18 of 21 High-dueling individuals were assigned to the gamergate cluster. Three High-dueling individuals exhibited intermediate phenotypes, including a single yolky oocyte and marginal hormonal levels. All Dropout and Low-dueling individuals are assigned to worker.

### Nutrition control

To assess the effects of nutritional state, four colonies were maintained under identical conditions but subjected to different feeding regimes: two calorie-restricted colonies (fed once on day 5) and two ad libitum colonies (fed on days 1, 3, and 5). On day 6, individuals from both conditions were combined to form four mixed colonies, which were recorded until day 24. Prior to mixing, four individuals died and were replaced to maintain equal colony sizes. After mixing, eight individuals died and were not replaced. Following 24 days of recording, individuals were collected for physiological and molecular analysis. Ovaries were dissected to quantify yolky oocyte number, and brain tissue was collected for measurement of corazonin (Cz) and vitellogenin (Vg) expression using quantitative PCR (qPCR), with expression levels normalized to internal housekeeping genes. Among the remaining individuals, 18 of 23 ad libitum ants were assigned to the gamergate caste, compared to 7 of 24 calorie-restricted ants. Differences in engaged-state occupancy between feeding conditions were assessed using a Mann-Whitney U test on per-individual disengaged-state occupancy.

### Trajectory prediction and Elo scores

Supervised classifiers were trained to distinguish prospective gamergates from workers using elastic-net logistic regression (scikit-learn). For each focal day, cumulative feature matrices were generated by summing daily values from day 1. Four predictor sets were compared: (1) an earlier “old quant” windowed measure of dueling activity, (2) the seven-feature daily dueling set, (3) the seven-feature set plus cumulative Elo drop-only variables derived from biting interactions, and (4) cumulative raw Elo-score variables alone.

Previous manual quantification proxy: This measure was based on sixteen 45-minute windows across the 12-hour observation period. An individual was scored as positive if it showed at least one initiative duel during the first 20 minutes of a window. Biting-derived Elo scores: Daily Elo ratings (initial Elo = 1000, K = 32) were updated chronologically for aggressive grooming and mandible locking events. The drop-only variable was defined as the decrease in Elo, where only losses were retained. Prediction evaluation used a nested leave-one-colony-out (LOCO) cross-validation framework, with performance quantified by the ROC AUC.

### Head width measurement

Head width (HW) was quantified from video recordings captured with a Basler acA2000 camera (2000 × 2000 pixel resolution) at full native resolution, ensuring maximal spatial precision without downsampling. For each individual, representative frames were extracted from tracking videos in which the head was clearly visible and oriented approximately perpendicular to the imaging plane. Prior to measurement, each frame was normalised for brightness uniformity and a Sobel gradient filter was applied in ImageJ to enhance edge contrast of the head capsule, improving landmark visibility. HW was defined as the maximum trans-orbital width: the straight-line distance between the outer lateral margins of the compound eyes, measured in full-face view. This corresponds to the widest point of the cranium and represents the standard morphometric landmark used in formicid body-size analyses. To ensure measurement accuracy, each line segment was constrained to be strictly perpendicular to the longitudinal midline axis of the head capsule, verified by overlaying a reference axis aligned to the clypeus–occiput midline; measurements deviating more than 5° from orthogonality were excluded and repeated. To reduce inter-measurement noise, HW was measured independently eight times per individual, and the mean value was taken as the final estimate. Pixel measurements were converted to millimeters using a calibration factor derived from known spatial scaling in the recording setup. The mean HW and standard deviation across the eight measurements are reported in Table 1.

