## Supplemental information for "Layered social competition coordinates reproductive hierarchy formation in ants"

**Supplementary Information**

**Supplementary Table S1. Morphological, physiological, and molecular measurements per individual**

| Colony | Ant_ID | Caste | Mean_HW_mm | Std_HW_mm | Ovaries | Corazonin | Vitellogenin |
|---|---|---|---|---|---|---|---|
| Col1 | 0 | dropout_worker | 2.044 | 0.0282 | 0 | 0.4217689431 | 0.0003993425329 |
| Col1 | 1 | low_dueling_worker | 2.085 | 0.0447 | 0 | 0.3775474358 | 0.00415667015 |
| Col1 | 2 | high_dueling_gamergate | 2.151 | 0.0371 | 3 | 0.05132399364 | 0.5033400178 |
| Col1 | 3 | dropout_worker | 2.044 | 0.0429 | 1 | 0.2441460213 | 0.003360795229 |
| Col1 | 4 | high_dueling_gamergate | 2.185 | 0.0493 | 3 | 0.0596884457 | 0.5033400178 |
| Col1 | 5 | low_dueling_worker | 2.079 | 0.0482 | 0 | 0.3522862301 | 0.0001603603477 |
| Col1 | 6 | dropout_worker | 1.990 | 0.0365 | 0 | 0.3301144009 | 0.06249932518 |
| Col1 | 7 | dropout_worker | 2.017 | 0.0176 | 0 | 0.3034169773 | 0.000265984686 |
| Col1 | 8 | high_dueling_gamergate | 2.121 | 0.0132 | 1 | 0.03056081141 | 0.09200012541 |
| Col1 | 9 | high_dueling_gamergate | 2.101 | 0.0489 | 1 | 0.05434476259 | 0.06699110607 |
| Col1 | 10 | high_dueling_gamergate | 2.206 | 0.0478 | 2 | 0.05515038131 | 0.04292976311 |
| Col1 | 11 | dropout_worker | 1.946 | 0.0123 | 0 | 0.2985686129 | 0.005210878747 |
| Col1 | 12 | low_dueling_worker | 1.977 | 0.0126 | 0 | 0.5912172302 | 0.001142554768 |
| Col1 | 13 | high_dueling_gamergate | 2.057 | 0.0245 | 4 | 0.08020757076 | 0.08774690306 |
| Col1 | 14 | high_dueling_gamergate | 1.955 | 0.0263 | 6 | 0.05870053789 | 1.040966992 |
| Col2 | 0 | high_dueling_gamergate | 2.054 | 0.066 | 5 | 0.08648017475 | 0.0005073007007 |
| Col2 | 1 | low_dueling_worker | 2.017 | 0.000 | NA | NA | NA |
| Col2 | 2 | dropout_worker | 2.030 | 0.003 | NA | NA | NA |
| Col2 | 3 | low_dueling_worker | 1.986 | 0.0343 | 0 | 0.5609257785 | 0.03027684877 |
| Col2 | 4 | low_dueling_worker | 1.997 | 0.0182 | NA | NA | NA |
| Col2 | 5 | high_dueling_gamergate | 1.928 | 0.0061 | 4 | 0.06295814277 | 0.03936333054 |
| Col2 | 6 | high_dueling_gamergate | 2.083 | 0.0146 | 1 | 0.0912165343 | 0.006385567293 |
| Col2 | 7 | low_dueling_worker | 2.035 | 0.0411 | 0 | 0.08998248051 | 0.0009009814053 |
| Col2 | 8 | dropout_worker | 2.089 | 0.0394 | NA | NA | NA |
| Col2 | 9 | high_dueling_gamergate | 1.993 | 0.0224 | 4 | 0.04767840038 | 0.1336468881 |
| Col2 | 10 | dropout_worker | 1.992 | 0.0197 | 1 | 1.010799608 | 0.03467267356 |
| Col2 | 11 | dropout_worker | 1.995 | 0.0328 | 0 | 0.523059244 | 0.0009788248392 |
| Col2 | 12 | high_dueling_gamergate | 2.098 | 0.0403 | 3 | 0.06933516876 | 0.09500579366 |
| Col2 | 13 | dropout_worker | 2.039 | 0.0388 | NA | NA | NA |
| Col2 | 14 | low_dueling_worker | 2.005 | 0.0174 | 0 | 0.02711721552 | 0.00117673146 |
| Col3 | 0 | low_dueling_worker | 1.949 | 0.0187 | 0 | 0.6426909565 | 0.000261244572 |
| Col3 | 1 | low_dueling_worker | 2.078 | 0.0279 | 1.5 | 0.2066352961 | 0.0462750107 |
| Col3 | 2 | high_dueling_gamergate | 2.127 | 0.0587 | 2.5 | 0.06063903258 | 0.1230869032 |
| Col3 | 3 | low_dueling_worker | 2.020 | 0.0141 | 0 | 0.1148724204 | 0.0001411201084 |
| Col3 | 4 | high_dueling_gamergate | 2.104 | 0.0374 | 2.5 | 0.009170108267 | 0.01698205848 |
| Col3 | 5 | low_dueling_worker | 1.946 | 0.0256 | 0 | 0.728678646 | 0.000857679982 |
| Col3 | 6 | high_dueling_gamergate | 2.074 | 0.0605 | 5 | 0.06383063331 | 0.0425140799 |
| Col3 | 7 | low_dueling_worker | 1.994 | 0.0167 | 1 | 0.2674793151 | 0.03286541435 |
| Col3 | 8 | low_dueling_worker | 2.051 | 0.0259 | 0 | 0.7131734612 | 0.00162442611 |
| Col3 | 9 | low_dueling_worker | 2.091 | 0.0190 | 1 | 0.3428907825 | 0.005030483593 |
| Col3 | 10 | high_dueling_gamergate | 2.048 | 0.0398 | 4 | 0.05866905075 | 0.0823710791 |
| Col3 | 11 | low_dueling_worker | 2.043 | 0.0260 | 0 | 0.2052262809 | 0.004482456404 |
| Col3 | 12 | low_dueling_worker | 1.919 | 0.0152 | 0 | 0.5789033472 | 0.004093193449 |
| Col3 | 13 | dropout_worker | 2.039 | 0.0491 | 0 | 0.07880526962 | 0.003467609636 |
| Col3 | 14 | high_dueling_gamergate | 2.094 | 0.0138 | NA | NA | NA |
| Col4 | 0 | dropout_worker | 2.092 | 0.0426 | 1 | NA | NA |
| Col4 | 1 | low_dueling_worker | 2.083 | 0.0932 | 0 | NA | NA |
| Col4 | 2 | high_dueling_gamergate | 2.106 | 0.0294 | 4 | NA | NA |
| Col4 | 3 | dropout_worker | 2.074 | 0.0199 | 0 | NA | NA |
| Col4 | 4 | low_dueling_worker | 2.105 | 0.0592 | 0 | NA | NA |
| Col4 | 5 | high_dueling_gamergate | 2.094 | 0.0599 | 2 | NA | NA |
| Col4 | 6 | high_dueling_gamergate | 2.070 | 0.0274 | 3 | NA | NA |
| Col4 | 7 | low_dueling_worker | 2.100 | 0.0246 | 0 | NA | NA |
| Col4 | 8 | dropout_worker | 2.012 | 0.0541 | 1 | NA | NA |
| Col4 | 9 | low_dueling_worker | 2.123 | 0.0377 | 0 | NA | NA |
| Col4 | 10 | dropout_worker | 2.066 | 0.0598 | 0 | NA | NA |
| Col4 | 11 | dropout_worker | 2.129 | 0.0734 | 1 | NA | NA |
| Col4 | 12 | low_dueling_worker | 2.095 | 0.0719 | 0 | NA | NA |
| Col4 | 13 | high_dueling_gamergate | 2.134 | 0.0499 | 2 | NA | NA |
| Col4 | 14 | low_dueling_worker | 2.116 | 0.0647 | 0 | NA | NA |

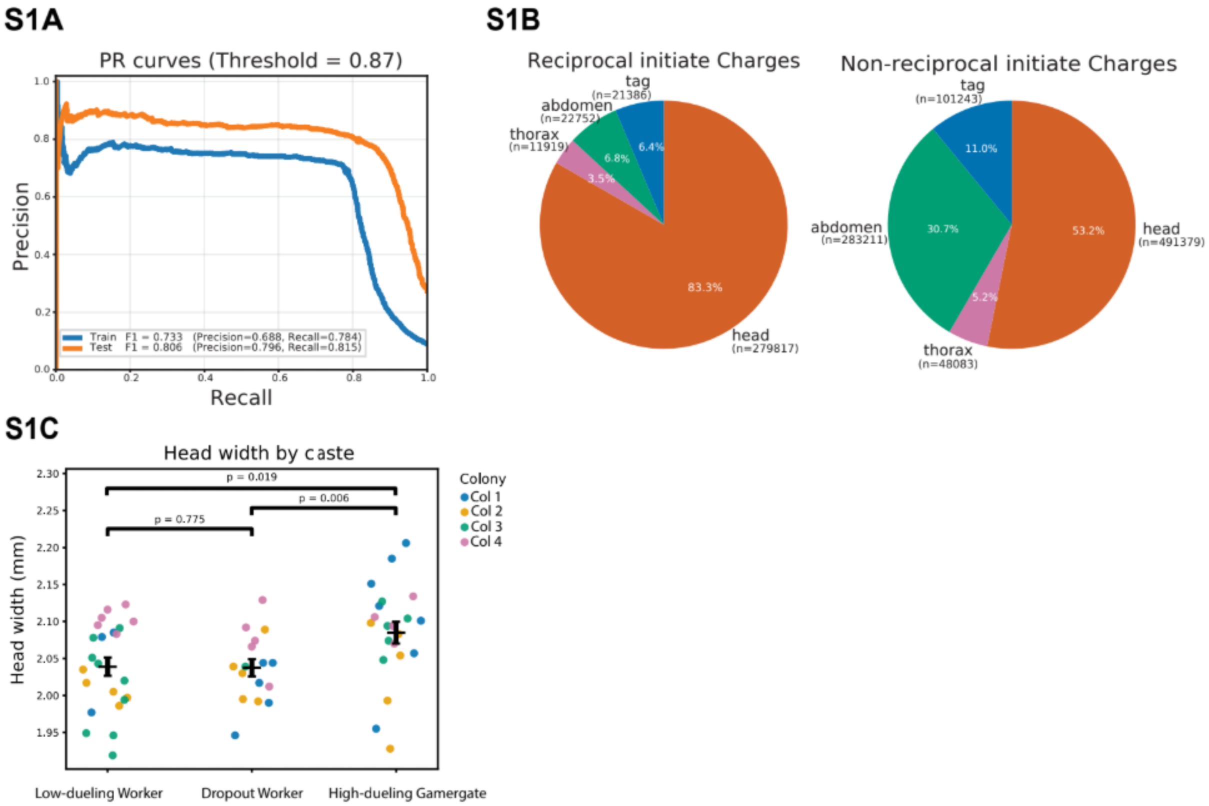


**Figure S1. Behavioral classifier performance and interaction composition. (A)** Precision–recall curves for the behavioral classifier distinguishing antennal interactions, showing strong performance on both training (F1 = 0.733) and test datasets (F1 = 0.806).

**(B)** Distribution of body-part contacts during interaction events, separated into reciprocal and non-reciprocal initiation.

**(C)** Head width comparison by dueling trajectory. P values were calculated using colony-blocked permutation tests.

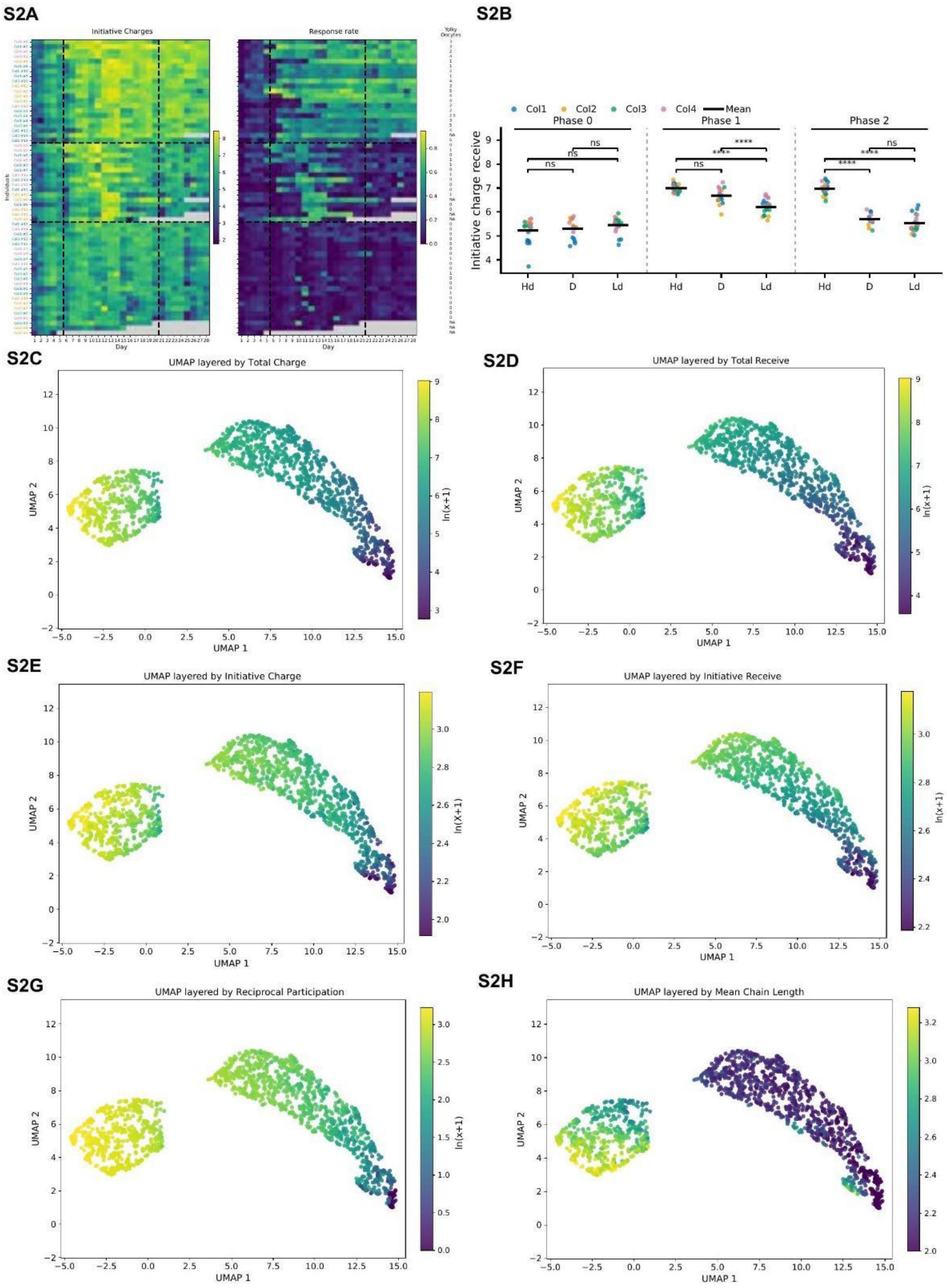

S2A
Initiative Charges
Response rate
Yolky Oocytes
Individuals
Day
S2B
Col1
Col2
Col3
Col4
Mean
Phase 0
Phase 1
Phase 2
Initiative charge receive
ns
****
Hd
D
Ld
S2C
UMAP layered by Total Charge
S2D
UMAP layered by Total Receive
S2E
UMAP layered by Initiative Charge
S2F
UMAP layered by Initiative Receive
S2G
UMAP layered by Reciprocal Participation
S2H
UMAP layered by Mean Chain Length
UMAP 1
UMAP 2
ln(x+1)

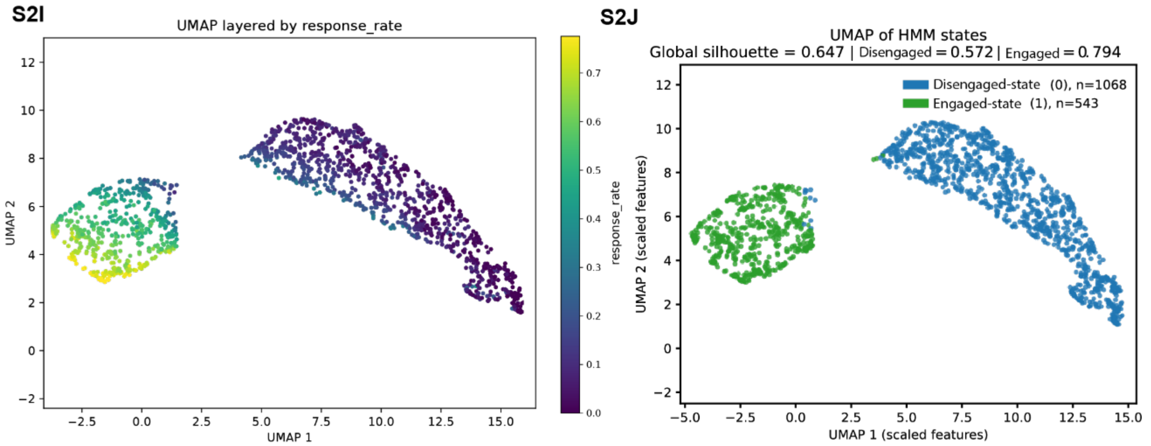


**Figure S2. Behavioral state structure and multidimensional embedding. (A)** Heatmaps of individual behavioral activity across days, showing structured temporal progression during caste transition. **(B)** Quantification of initiative charge received across behavioral classes (High-dueling, Dropout, Low-dueling) and phases, revealing significant divergence during engagement and stabilization phases. **(C-I)** UMAP embeddings of behavioral features, colored by total charge, total receive, initiative charge, initiative receive, reciprocal participation, mean interaction chain length, and response rate respectively. These representations reveal a continuous behavioral manifold that separates into distinct regions corresponding to behavioral states. **(J)** Hidden Markov Model (HMM) state assignment projected onto the UMAP embedding. Two discrete states are identified, corresponding to an engaged state and a disengaged state, with strong separation (global silhouette = 0.647; disengaged-state mean = 0.572; engaged-state mean = 0.794). The two states occupy distinct regions of the behavioral manifold, indicating that continuous variation in interaction features organizes into discrete, stable behavioral states.

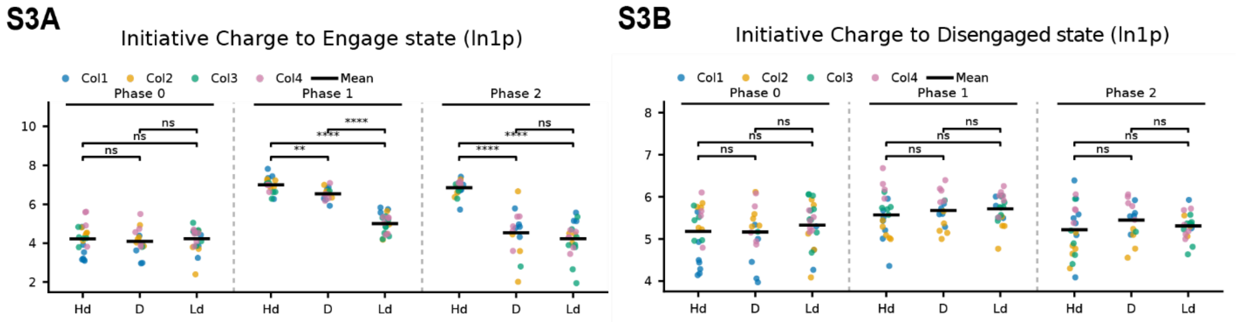


**Figure S3. State-dependent interaction directionality. (A)** Initiative charges directed toward engaged-state individuals across phases, showing increased targeting during engagement and stabilization phases, with significant differences between behavioral classes.
**(B)** Initiative charge directed toward disengaged-state individuals, showing no consistent differences across classes or phases. These results indicate that interaction asymmetry is selectively enhanced toward engaged-state individuals, supporting preferential interaction among reproductively active ants.

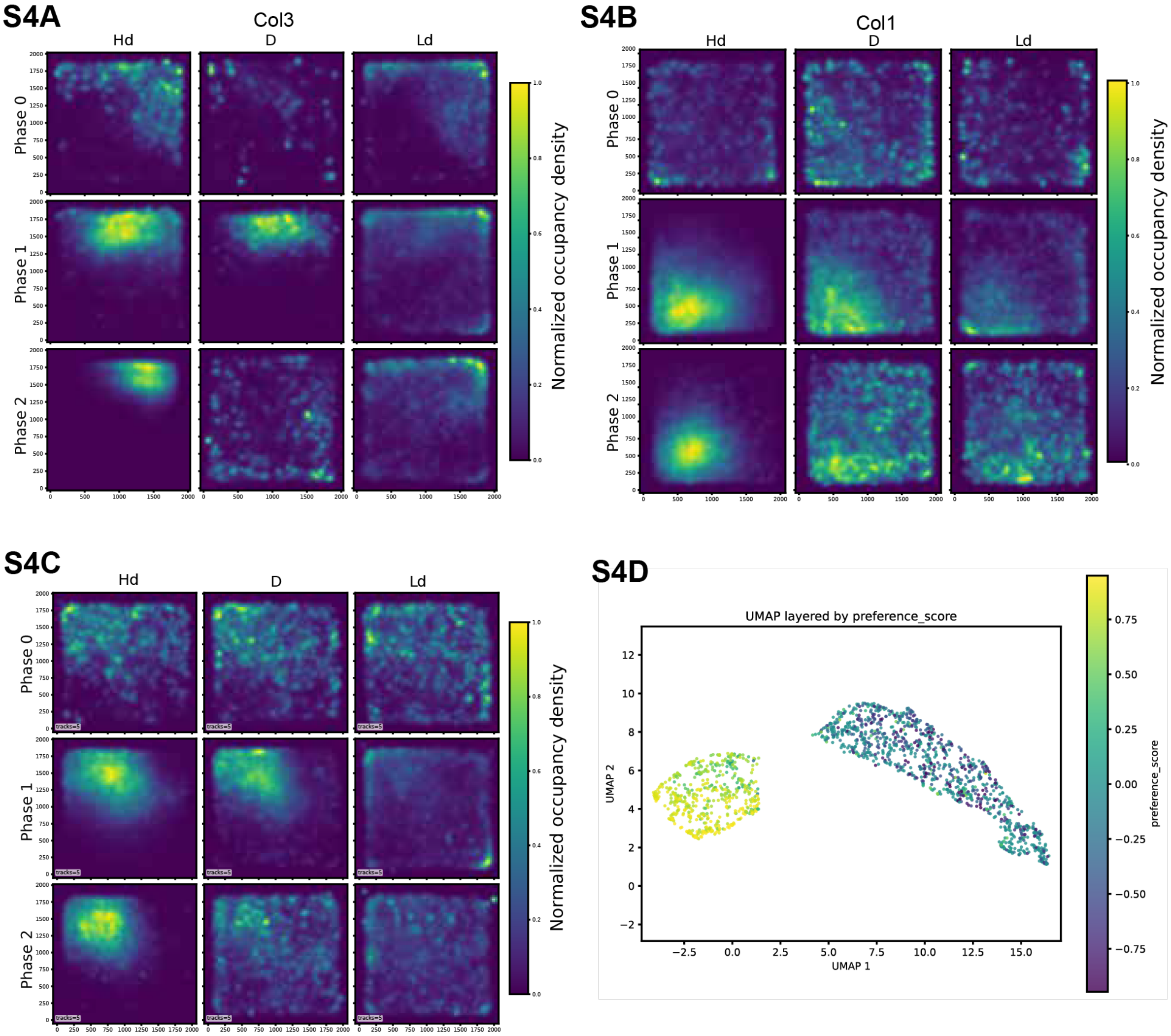


**Figure S4. Spatial occupancy dynamics across behavioral states. (A-C)** Spatial occupancy density maps across phases for representative colonies, separated by behavioral class (High-dueling, Dropout, Low-dueling). Engaged-state individuals progressively concentrate within a restricted spatial region, forming a centralized interaction zone. These results demonstrate that behavioral states are associated with distinct spatial organization within the colony. **(D)** UMAP embeddings of preference score. These representations reveal a continuous behavioral manifold that separates into distinct regions corresponding to behavioral states.

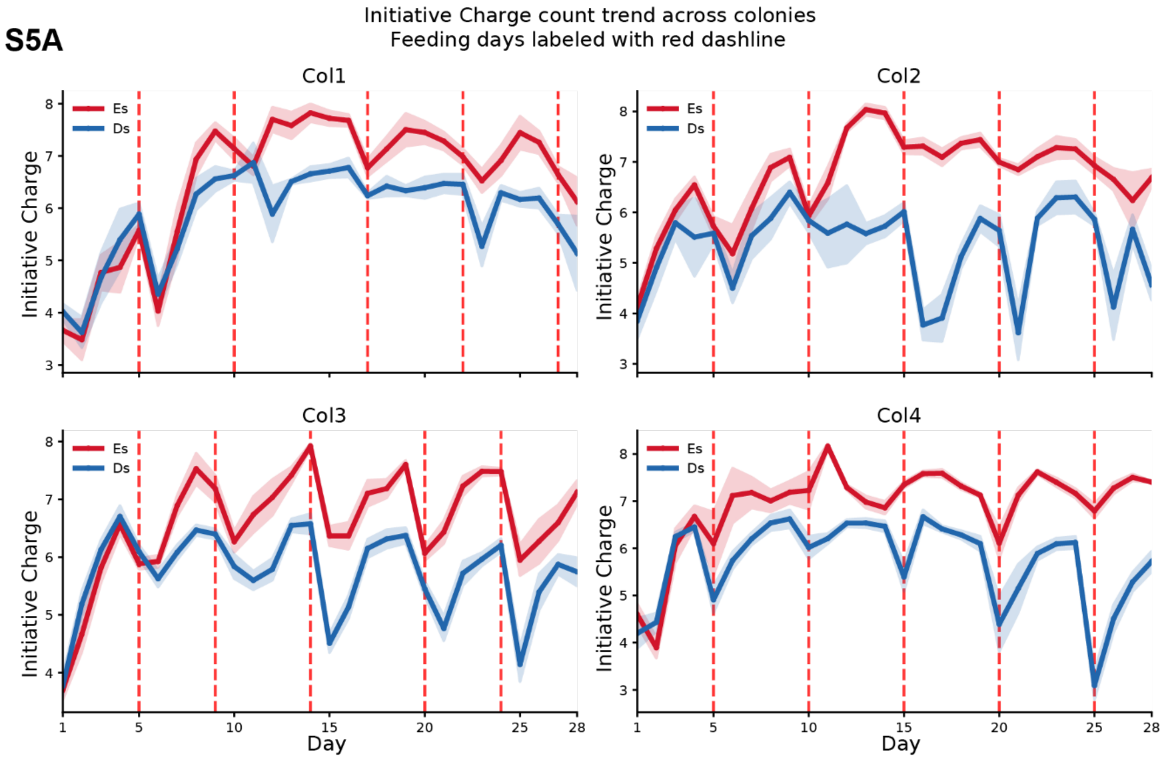


**Figure S5. Temporal dynamics of initiative charge and feeding schedule across colonies. (A)** Initiative charge counts over time for each colony, separated into engaged-state and disengaged-state individuals. Red dashed lines indicate feeding events. Engaged-state individuals (Es) exhibit sustained elevated interaction levels, whereas disengaged-state individuals (Ds) show reduced and more variable activity. Temporal fluctuations align with feeding events but preserve overall state-dependent differences.

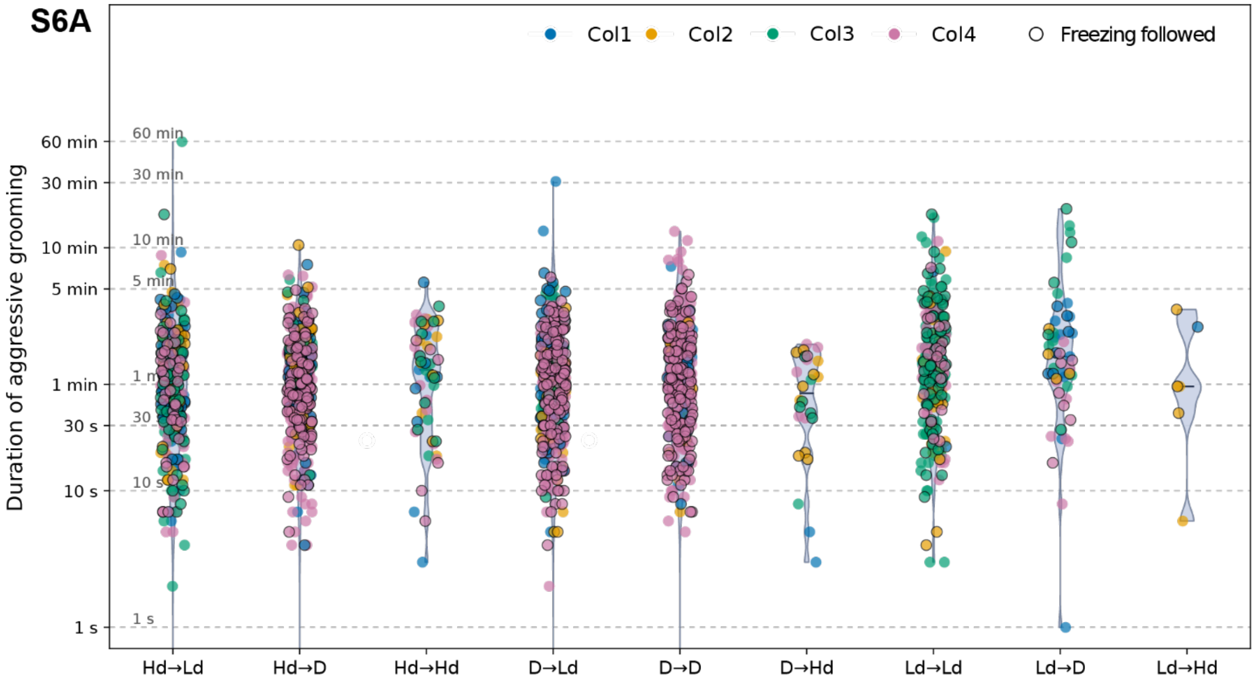
S6A
Col1
Col2
Col3
Col4
Freezing followed
Duration of aggressive grooming
60 min
30 min
10 min
5 min
1 min
30 s
10 s
1 s
Hd→Ld
Hd→D
Hd→Hd
D→Ld
D→D
D→Hd
Ld→Ld
Ld→D
Ld→Hd

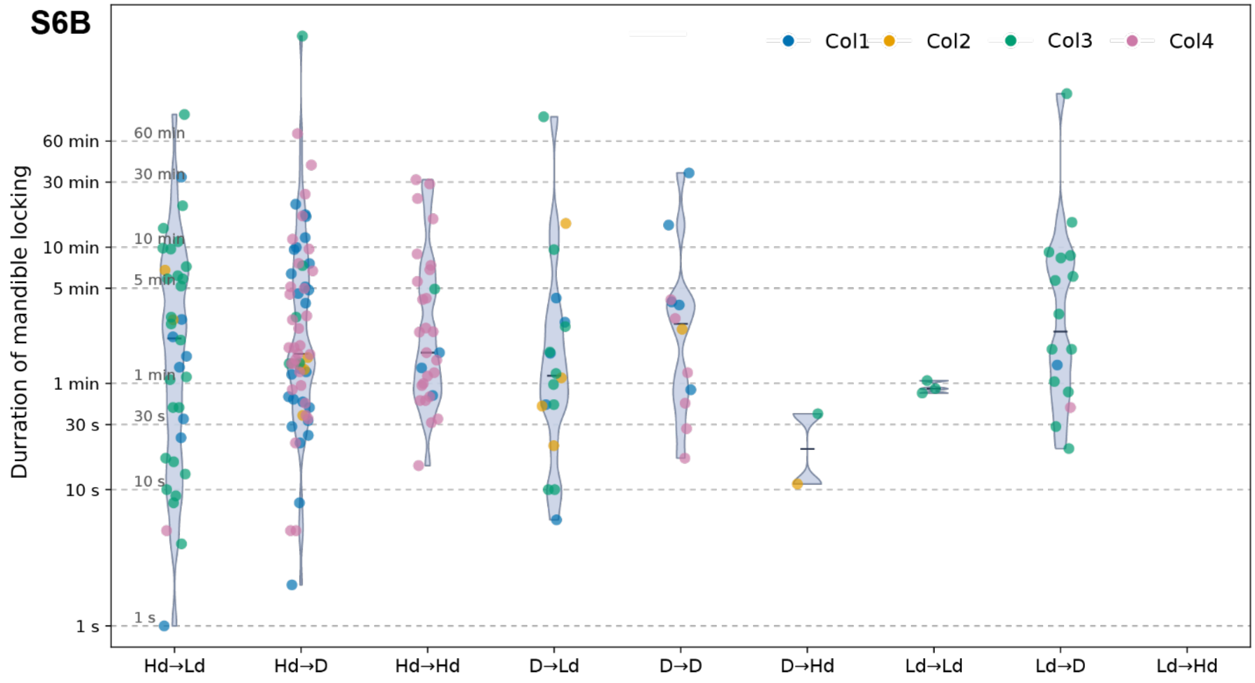
S6B
Col1
Col2
Col3
Col4
Durration of mandible locking
60 min
30 min
10 min
5 min
1 min
30 s
10 s
1 s
Hd→Ld
Hd→D
Hd→Hd
D→Ld
D→D
D→Hd
Ld→Ld
Ld→D
Ld→Hd

64

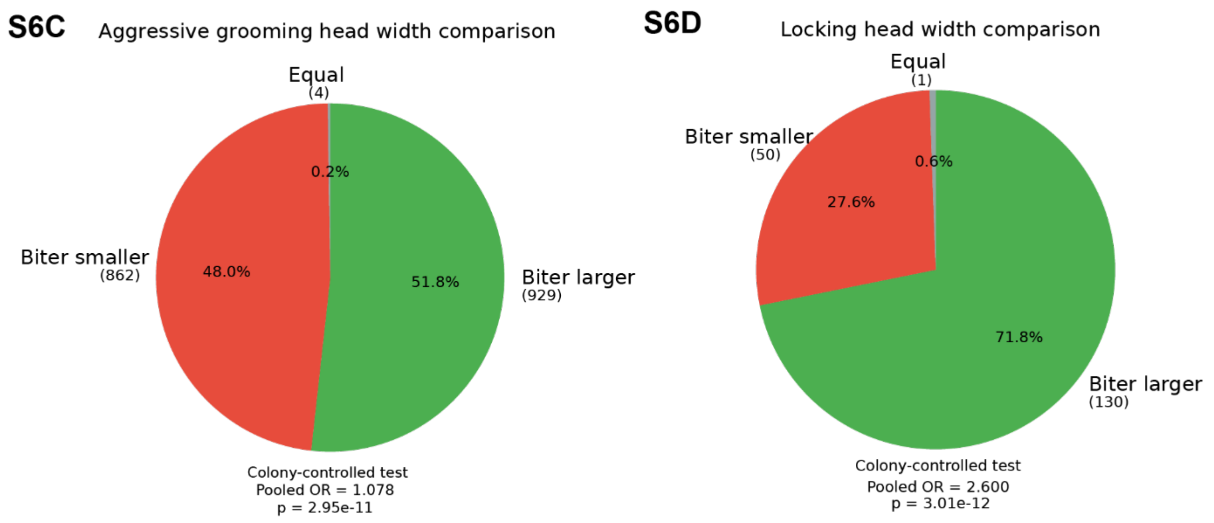


**Figure S6. Physical interaction outcomes and size asymmetry**. **(A)** Distribution of aggressive grooming durations across interaction types, showing prolonged interactions in specific pairings. **(B)** Duration of mandible locking events, indicating distinct temporal signatures compared to grooming. **(C)** Head width comparison for aggressive grooming interactions, showing no strong bias toward larger individuals. **(D)** Head width comparison for mandible locking events, revealing a significant bias toward larger individuals initiating interactions.
These results indicate that different interaction types exhibit distinct physical and size-dependent asymmetries.
